# VIP plasma levels associate with survival in severe COVID-19 patients, correlating with protective effects in SARS-CoV-2-infected cells

**DOI:** 10.1101/2020.07.25.220806

**Authors:** Jairo R. Temerozo, Carolina Q. Sacramento, Natalia Fintelman-Rodrigues, Camila R. R. Pão, Caroline S. de Freitas, Suelen Silva Gomes Dias, André C. Ferreira, Mayara Mattos, Vinicius Cardoso Soares, Lívia Teixeira, Isaclaudia G. Azevedo-Quintanilha, Eugenio D. Hottz, Pedro Kurtz, Fernando A. Bozza, Patrícia T. Bozza, Thiago Moreno L. Souza, Dumith Chequer Bou-Habib

## Abstract

Infection by SARS-CoV-2 may elicit uncontrolled and damaging inflammatory responses. Thus, it is critical to identify compounds able to inhibit virus replication and thwart the inflammatory reaction. Here, we show that the plasma levels of the immunoregulatory neuropeptide VIP are elevated in patients with severe COVID-19, correlating with reduced inflammatory mediators and with survival on those patients. In vitro, VIP and PACAP, highly similar neuropeptides, decreased the SARS-CoV-2 genome replication in human monocytes and viral production in lung epithelial cells, also reducing cell death. Both neuropeptides inhibited the production of proinflammatory mediators in lung epithelial cells and in monocytes. VIP and PACAP prevented in monocytes the SARS-CoV-2-induced activation of NF-kB and SREBP1 and SREBP2, transcriptions factors involved in proinflammatory reactions and lipid metabolism, respectively. They also promoted CREB activation, a transcription factor with antiapoptotic activity and negative regulator of NF-kB. Specific inhibition of NF-kB and SREBP1/2 reproduced the anti-inflammatory, antiviral and cell death protection effects of VIP and PACAP. Our results support further clinical investigations of these neuropeptides against COVID-19.

## Introduction

Individuals with coronavirus disease 2019 (COVID-19), caused by the severe acute respiratory syndrome coronavirus 2 (SARS-CoV-2) [1], may present asymptomatic or mild disease to severe lung inflammation and acute respiratory distress syndrome (ARDS) [2,3], besides a variety of extrapulmonary manifestations [4]. Severe SARS-CoV-2 infection is characterized by elevated serum levels of proinflammatory mediators (hypercytokinemia, also known as cytokine storm) such as, for example, IL-2, IL-6, TNF, IL-8, IL-1β, IFN-γ [2,3,5,6]. The dysregulated immune response and production of cytokines and chemokines are hallmarks of SARS-CoV-2 infection and have been pointed as the main cause of the severe lung damage and unfavorable clinical progression of patients with COVID-19 [3–8]. Also, the in vivo formation of neutrophil extracellular traps (NETs) in the lungs, SARS-CoV-2-induced inflammasome activation and cell death by pyroptosis, have also been considered as risk factors in critically ill COVID-19 patients [9–14].

During the inflammatory response to human pathogenic coronaviruses, circulating neutrophils and monocytes migrate and infiltrate the lungs [15,16] and other organs, contributing to potentiate and perpetuate the inflammation and eventually exacerbating the tissue damage [17–19]. Previous studies showed that MERS-CoV- and SARS-CoV-infected macrophages produce high levels of pro-inflammatory cytokines and chemokines [20,21], and, more recently, that lung monocytes from severe pneumonia caused by SARS-CoV-2 are potent producers of TNF-α and IL-6, whose levels were increased in the serum of the same patients [7]. Also, we and other authors have found that SARS-CoV-2 induces inflammasome activation and cell death by pyroptosis in monocytes, either by experimental or natural infection, which are associated with lung inflammation and are risk factors in critically ill COVID-19 patients [13,14].

Thus, it is critical to identify agents able to prevent the infection and concurrently thwart the prototypical dysregulated inflammatory reaction and tissue lesions secondary to SARS-CoV-2 infection. In this work, we evaluated whether the neuropeptides Vasoactive Intestinal Peptide (VIP) and Pituitary Adenylate Cyclase-Activating Polypeptide (PACAP) can present protective effects in SARS-CoV-2 infection. VIP and PACAP share many biological properties through their interaction with the G protein-coupled receptors VPAC1, VPAC2 and PAC1 [22], which are systemically distributed. They have well-characterized regulatory effects on the immune system and anti-inflammatory properties, including control of cell activation and differentiation, down-regulation of inflammatory cytokines and reactive oxygen species and induction of the anti-inflammatory cytokine IL-10 [23–28]. Based on their consistent anti-inflammatory and pro-homeostatic activities, both neuropeptides have been considered as promising therapeutic agents for autoimmune disorders and chronic inflammatory illnesses [29–32]. Therefore, based on the well-known properties of both neuropeptides to regulate inflammatory reactions, and on the dysregulated immune responses that affect COVID-19 patients, we investigated whether they could present protective roles during SARS-CoV-2 infection. We report here that VIP levels are elevated in the plasma of individuals with severe manifestations of COVID-19, which correlated with survival on critically ill patients. We also verified, in in vitro assays, that VIP and PACAP inhibit the production of proinflammatory mediators in SARS-CoV-2-infected monocytes and lung epithelial cells; and reduced viral production and cell death.

## Materials and Methods

### Cells, virus and reagents

African green monkey kidney cells (Vero, subtype E6) and human lung epithelial cell lines (Calu-3) were expanded in high glucose DMEM (Vero) or MEM (Calu-3) with 10% fetal bovine serum (FBS; Merck), with 100 U/mL penicillin and 100 μg/mL streptomycin (Pen/Strep; Gibco) at 37°C in a humidified atmosphere with 5% CO_2_. Peripheral blood mononuclear cells (PBMCs) were isolated by density gradient centrifugation (Ficoll-Paque, GE Healthcare) from buffy-coat preparations of blood from healthy donors. PBMCs (2 × 10^6^ cells) were plated onto 48-well plates (NalgeNunc) in RPMI-1640 with 5% inactivated male human AB serum (Merck) for 3 hours. Non-adherent cells were removed and monocytes were maintained in DMEM (low-glucose) with 5% human serum and 100 U/mL penicillin and 100 μg/mL streptomycin. Purity of monocytes was above 90%, as determined by flow cytometry (FACScan; Becton Dickinson) using anti-CD3 (BD Biosciences) and anti-CD14 (BD Biosciences) antibodies. SARS-CoV-2 (GenBank accession no. MT710714) was expanded in Vero E6 cells. Viral isolation was performed after a single passage in a cell culture in a 150 cm^2^ flasks with high glucose DMEM plus 2% FBS. Observations for cytopathic effects were performed daily and peaked 4 to 5 days after infection. All procedures related to virus culture were handled in biosafety level 3 (BSL3) multiuser facilities, according to WHO guidelines. Virus titers were determined as plaque forming units (PFU/mL), and virus stocks were kept in −80°C ultralow freezers. VIP and PACAP and the VPAC1 and VPAC2 agonists (Ala^11,22,28^)-VIP and Bay 55-9837, respectively, were purchased from Tocris. The PAC1 agonist Maxadilan was kindly donated by Dr. Ethan A. Lerner (Department of Dermatology, Massachusetts General Hospital, MA, USA). All peptides and agonists were diluted in PBS. The inhibitors of the transcription factors SREBP (AM580) and NF-KB (Bay 11-7082) were purchased from Selleckchem.

### Infections and virus titration

Infections were performed with SARS-CoV-2 at MOI of 0.01 (monocytes) or 0.1 (Calu-3) in low (monocytes) or high (Calu-3) glucose DMEM without serum. After 1 hour, viral input was removed and cells were washed and incubated with complete medium with treatments or not. For virus titration, monolayers of Vero E6 cells (2 × 10^4^ cell/well) in 96-well plates were infected with serial dilutions of supernatants containing SARS-CoV-2 for 1 hour at 37°C. Semi-solid high glucose DMEM medium containing 2% FBS and 2.4% carboxymethylcellulose was added and cultures were incubated for 3 days at 37 °C. Then, the cells were fixed with 10% formalin for 2 hours at room temperature. The cell monolayer was stained with 0.04% solution of crystal violet in 20% ethanol for 1 hour. Plaque numbers were scored in at least 3 replicates per dilution by independent readers blinded to the experimental group, and the virus titers were determined by plaque-forming units (PFU) per milliliter.

### Molecular detection of virus RNA levels

The total RNA was extracted from cells using QIAamp Viral RNA (Qiagen), according to manufacturer’s instructions. Quantitative RT-PCR was performed using QuantiTect Probe RT-PCR Kit (Qiagen) in a StepOnePlus™ Real-Time PCR System (Thermo Fisher Scientific). Amplifications were carried out in 15 µL reaction mixtures containing 2× reaction mix buffer, 50 µM of each primer, 10 µM of probe, and 5 µL of RNA template. Primers, probes, and cycling conditions recommended by the Centers for Disease Control and Prevention (CDC) protocol were used to detect the SARS-CoV-2 [33]. The standard curve method was employed for virus quantification. For reference to the cell amounts used, the housekeeping gene RNAse P was amplified. The Ct values for this target were compared to those obtained to different cell amounts, 10^7^ to 10^2^, for calibration.

### SDS-PAGE and Western blot for SREBPs

After 24h of SARS-CoV-2 infection, monocytes were harvested using ice-cold lysis buffer (1% Triton X-100, 2% SDS, 150 mM NaCl, 10 mM HEPES, 2 mM EDTA containing protease inhibitor cocktail - Roche). Cell lysates were heated at 100 °C for 5 min in the presence of Laemmli buffer (20% β-mercaptoethanol; 370 mM Tris base; 160 μM bromophenol blue; 6% glycerol; 16% SDS; pH 6.8), and 20 μg of protein/sample were resolved by electrophoresis on SDS-containing 10% polyacrylamide gel (SDS-PAGE). After electrophoresis, the separated proteins were transferred to nitrocellulose membranes and incubated in blocking buffer (5% nonfat milk, 50 mM Tris-HCl, 150 mM NaCl, and 0.1% Tween 20). Membranes were probed overnight with the following antibodies: anti-SREBP-1 (Proteintech #14088-1-AP), anti-SREBP-2 (Proteintech #28212-1-AP) and anti-β-actin (Sigma, #A1978). After the washing steps, they were incubated with IRDye - LICOR or HRP-conjugated secondary antibodies. All antibodies were diluted in blocking buffer. The detections were performed by Supersignal Chemiluminescence (GE Healthcare) or by fluorescence imaging using the Odyssey system. Densitometries were analyzed using the Image Studio Lite Version 5.2 software.

### Measurements of inflammatory mediators, cell death, NF-kBp65, CREB and neuropeptides

A multiplex biometric immunoassay containing fluorescent dyed microbeads was used to measure cytokines in plasma samples (Bio-Rad Laboratories). The following cytokines were quantified: Basic-FGF, CTRAK, Eotaxin, G-CSF, GRO-α, HGF, IFN-α2, IFN-β, IFN-γ, IL-1α, IL-1β, IL-1RA, IL-2, IL-2RA, IL-3, IL-4, IL-5, IL-6, IL-7, IL-8, IL-9, IP-10, IL-10, IL-12(p40), IL-13, IL-15, IL-16, IL-17A, IL-18, LIF, M-CSF, MCP-3, MIF, MIG, MIP-1β, PDGF-BB, RANTES, SCF, SCGF-1α, SCGF-β, TNFα, TNFβ, VEGF, β-NGF and PF4. Cytokine levels were calculated by Luminex technology (Bio-Plex Workstation; Bio-Rad Laboratories). The analysis of data was performed using software provided by the manufacturer (Bio-Rad Laboratories). A range of 0.51–8000 pg/mL recombinant cytokines was used to establish standard curves and the sensitivity of the assay. The levels of IL-6, IL-8, TNF-α and MIF were quantified in the supernatants from uninfected and SARS-CoV-2-infected Calu-3 cells and monocytes by ELISA (R&D Systems), following manufacturer’s instructions, and results are expressed as percentages relative to uninfected cells. Cell death was determined according to the activity of lactate dehydrogenase (LDH) in supernatants using CytoTox® Kit (Promega) according to the manufacturer’s instructions. Supernatants were centrifuged at 5,000 rpm for 1 minute to remove cellular debris. Evaluation of NF-kBp65 and CREB activation was performed in infected or uninfected monocytes using NF-kB p65 (Total/Phospho) InstantOne™ and CREB (Total/Phospho) Multispecies InstantOne™ ELISA Kits (Thermo Fisher), according to manufacturer’s instructions. VIP and PACAP levels were quantified in the plasma from patients or control volunteers using standard commercially available ELISA and EIA Kits, according to the manufacturer’s instructions (Abelisa).

### Human subjects

We prospectively enrolled patients with severe or mild/asymptomatic COVID-19 RT-PCR-confirmed diagnosis, and SARS-CoV-2-negative healthy controls. Blood and respiratory samples were obtained from 24 patients with severe COVID-19 within 72 hours from intensive care unit (ICU) admission in two reference centers (Instituto Estadual do Cérebro Paulo Niemeyer and Hospital Copa Star, Rio de Janeiro, Brazil). Severe COVID-19 was defined as those critically ill patients presenting viral pneumonia on computed tomography scan and requiring oxygen supplementation through either a nonrebreather mask or mechanical ventilation. Eight outpatients presenting mild self-limiting COVID-19 syndrome, and two SARS-CoV-2-positive asymptomatic subjects were also included. Patients had SARS-CoV-2 confirmed diagnostic through RT-PCR of nasal swab or tracheal aspirates. Peripheral vein blood was also collected from 10 SARS-CoV-2-negative healthy participants as tested by RT-PCR on the day of blood sampling. Characteristics of severe (n=24), mild/asymptomatic (n=10) and healthy (n=10) participants are presented in **Table 1**. Mild and severe COVID-19 patients presented differences regarding age and presence of comorbidities, such as obesity, cardiovascular diseases and diabetes (**Table 1**), which is consistent with previously reported patient cohorts [2,34–36]. The SARS-CoV-2-negative control group included subjects of older age and chronic non-communicable diseases, so it is matched with mild and critical COVID-19 patients, except for hypertension (**Table 1**). All ICU-admitted patients received usual supportive care for severe COVID-19 and respiratory support with either noninvasive oxygen supplementation (n=5) or mechanical ventilation (n=19) (**Supplemental Table 1**). Patients with acute respiratory distress syndrome (ARDS) were managed with neuromuscular blockade and a protective ventilation strategy that included low tidal volume (6 mL/kg of predicted body weight) and limited driving pressure (less than 16 cmH2O) as well as optimal PEEP calculated based on the best lung compliance and PaO2/FiO2 ratio. In those patients with severe ARDS and PaO2/FiO2 ratio below 150 despite optimal ventilatory settings, prone position was initiated. Our management protocol included antithrombotic prophylaxis with enoxaparin 40 to 60 mg per day. Patients did not receive routine steroids, antivirals or other anti-inflammatory or anti-platelet drugs. The SARS-CoV-2-negative control participants were not under anti-inflammatory or anti-platelet drugs for at least two weeks. All clinical information was prospectively collected using a standardized form - ISARIC/WHO Clinical Characterization Protocol for Severe Emerging Infections (CCPBR). Clinical and laboratory data were recorded on admission in all severe patients included in the study and the primary outcome analyzed was 28-day mortality (n = 11 survivors and 13 non-survivors, **Supplemental Table 2**). Age and frequency of comorbidities were not different between severe patients requiring mechanical ventilation or noninvasive oxygen supplementation neither between survivors and non-survivors (**Supplemental Table 1 and 2**).

**Table 1:** Characteristics of COVID-19 patients and control subjects.

| Characteristics<sup>1</sup> | Control<br>(n=10) | Asymptomatic/<br>Mild<br>(n=10) | Severe/critical<br>(n=24) |
| --- | --- | --- | --- |
| Age, years | 53 (32 – 60) | 43 (24 – 52) | 58 (48 – 66) |
| Sex, male | 4 (40%) | 4 (40%) | 12 (50%) |
| Respiratory support |  |  |  |
| Oxygen supplementation | 0 (0%) | 0 (0%) | 5 (20.8%) |
| Mechanical ventilation | 0 (0%) | 0 (0%) | 19 (79.2%) |
| SAPS 3 | - | - | 60 (55 – 71) |
| PaO<sub>2</sub>/FiO<sub>2</sub> ratio | - | - | 154 (99 – 373) |
| Vasopressor | - | - | 10 (41.6%) |
| Time from symptom onset<br>to blood sample, days | - | 6 (-1 – 8)<sup>2</sup> | 14 (8 – 17) |
| 28-day mortality | - | - | 13 (54.2%) |
| **Comorbidities** |  |  |  |
| Obesity | 1 (10%) | 1 (10%) | 5 (20.8%) |
| Hypertension | 1 (10%) | 2 (20%) | 6 (25%) |
| Diabetes | 0 (0%) | 0 (0%) | 9 (37.5%) |
| Cancer | 0 (0%) | 0 (0%) | 3 (12.5%) |
| Heart disease<sup>3</sup> | 0 (0%) | 0 (0%) | 2 (8.3%) |
| **Presenting symptoms** |  |  |  |
| Cough | 0 (0%) | 3 (30%) | 17 (70.8%) |
| Fever | 0 (0%) | 5 (50%) | 18 (75%) |
| Dyspnea | 0 (0%) | 0 (0%) | 20 (83.3%) |
| Headache | 0 (0%) | 4 (40%) | 3 (12.5%) |
| Anosmia | 0 (0%) | 4 (40%) | 8 (33.3%) |
| **Laboratory findings on admission** |  |  |  |
| Leukocytes, x 1000/μL | - | - | 138 (102 – 180) |
| Lymphocyte, cells/μL | - | - | 1,167 (645 – 1,590) |
| Monocytes, cells/μL | - | - | 679 (509 – 847) |
| Platelet count, x 1000/μL | - | - | 169 (137 – 218) |
| C Reactive Protein, mg/L<sup>4</sup> | 0.1 (0.1 – 0.18) | 0.2 (0.1 – 0.13) | 178 (74 – 308)* |
| Fibrinogen, mg/dL<sup>4</sup> | 281 (232 – 302) | 248 (182 – 341) | 528 (366 – 714)* |
| D-dimer, IU/mL<sup>4</sup> | 292 (225 – 476) | 191 (187 – 313) | 4836 (2364 – 10816)* |

<sup>1</sup>Numerical variables are represented as the median and the interquartile range, and qualitative variables are represented as the number and the percentage.
<sup>2</sup>Day of sample collection after the onset of symptoms was not computed for asymptomatic subjects.
<sup>3</sup>Coronary artery disease or congestive heart failure.
<sup>4</sup>Reference values of C reactive Protein (0.00 – 1.00), Fibrinogen (238 – 498 mg/dL) and D-dimer (0 – 500 ng/mL).
\*p < 0.05 compared to control. The qualitative variables were compared using the two tailed Fisher exact test, and the numerical variables using the t test for parametric and the Mann Whitney U test for nonparametric distributions.

### Statistical analysis

Statistics were performed using GraphPad Prism software version 8. Numerical variables were tested regarding distribution using the Shapiro-Wilk test. One-way analysis of variance (ANOVA) was used to compare differences among 3 groups following a normal (parametric) distribution, and Tukey’s post-hoc test was used to locate the differences between the groups; or Friedman’s test (for non-parametric data) with Dunn’s post-hoc test. Comparisons between 2 groups were performed using the Student t test for parametric distributions or the Mann-Whitney U test for nonparametric distributions. Correlation coefficients were calculated using Pearson’s correlation test for parametric distributions and the Spearman’s correlation test for nonparametric distributions.

## Results

### Plasma levels of VIP are elevated in patients with severe forms of COVID-19 and associate with survival

From April to May 2020, we followed up 24 critically ill COVID-19 patients, at the median age of 53-year-old (**Table 1**), presenting the most common infection symptoms and comorbidities, from whom we evaluated the plasma levels of the neuropeptides VIP and PACAP, comparing with patients with mild COVID-19 symptoms and non-infected healthy individuals. We found that patients affected by the most severe forms of infection had higher plasma levels of the neuropeptide VIP than uninfected healthy controls and asymptomatic/mild patients (**Fig. 1A**). Comparing the viral load in positive swab samples from mild and severe COVID-19 patients we found a modest positive correlation with VIP levels (**Fig. 1B**). Following, we examined a possible correlation between VIP levels of severe patients and inflammatory markers. We identified that VIP negatively correlated with five pro-inflammatory factors (IL-8, IL-12p40, IL-17A, TNF-α and CXCL10/IP-10), and positively with two anti-inflammatory factors (IL-1RA and IL-10) (**Fig. 1C-I**). Next, severe COVID-19 patients were further subdivided between those requiring invasive mechanical ventilation or noninvasive O_2_ supplementation or according to the 28-day mortality outcome as survivors or non-survivors. We did not find a significant difference when analyzing O2 supplementation versus mechanical ventilation (**Fig. 1J**), probably due to the low number of patients under the first condition. On the contrary, we observed that VIP plasma levels associated with survival of patients with severe COVID-19, being significantly higher in survivors than in non-survivors (**Fig. 1K**). For PACAP plasma levels, we did not find significant differences between the groups analyzed, inflammatory markers, viral load or with VIP levels (data not shown). The finding that survival of severe COVID-19 patients is associated with higher levels of circulating VIP, a molecule with pro-homeostasis and anti-inflammatory activities [32,37], moreover pointing to an application as a prognostic marker, also implies to a therapeutical potential of VIP in COVID-19. In fact, VIP has been approved for three clinical trials against COVID-19 in intravenous [38] and inhaled [39,40] formulations. Our initial clinical data prompted us to evaluate the effects of VIP (and of PACAP as well) on SARS-CoV-2-infected cells to better corroborate the use of VIP as therapeutical agent in COVID-19 patients.

**Figure 1.**
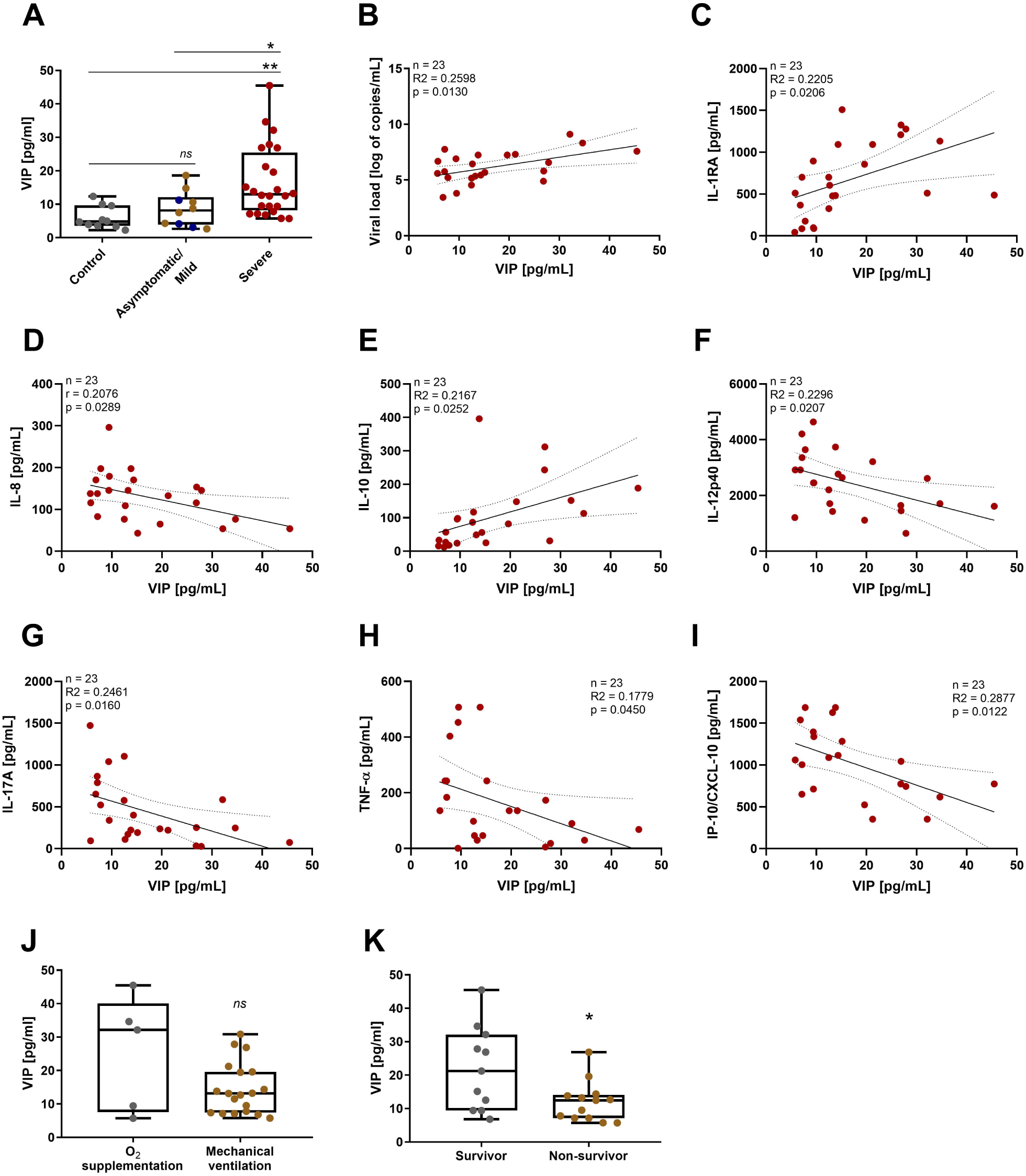
Plasma levels of VIP are elevated in patients with severe forms of COVID-19 and associates with reduced levels of inflammatory markers and with survival. The levels of VIP (**A**) in the plasma of SARS-CoV-2-negative control participants, SARS-CoV-2-positive asymptomatic subjects, or symptomatic patients presenting mild to severe COVID-19 were quantified by ELISA. Correlation between levels of VIP and viral load (**B**) or inflammation markers (**C-I**). Severe COVID-19 patients admitted to the ICU were sub-divided between those requiring invasive mechanical ventilation or noninvasive O_2_ supplementation (**J**) and according to the 28-day mortality outcome as survivors or non survivors (**K**). Linear regression (with the 95% confidence interval) and Spearman’s correlation were calculated according to the distribution of the data. Dots represent: Controls, grey; Asymptomatics, blue; Mild, brown; Severe, red). The horizontal lines in the box plots represent the median, the box edges represent the interquartile ranges, and the whiskers indicate the minimal and maximal value in each group. \**p* ≤ .05; \*\**p* ≤ .01.

### VIP and PACAP reduce SARS-CoV-2 RNA synthesis in human primary monocytes and viral replication in pulmonary cells, protecting them from virus-mediated cytopathic effects

Upon identifying the association of VIP with survival of critical COVID-19 patients and considering that, in the setting of COVID-19, the main affected cells are those present in the lung epithelium, including the immune cells recruited upon infection, we sought to investigate the in vitro effects of VIP and PACAP in SARS-CoV-2-infected cells. To this end, we initially evaluated the SARS-CoV-2 RNA synthesis in monocytes (as the infection by SARS-CoV-2 in this cell is non-productive [41,42]) and the viral replication in Calu-3 cells (a lineage of lung epithelial cells highly susceptible to SARS-CoV-2) exposed to VIP or PACAP). We found that VIP reduced the SARS-CoV-2 RNA synthesis in monocytes, achieving up to 40% and 50% inhibition at 5 nM and 10 nM, respectively (**Fig. 2A**). PACAP similarly decreased the levels of viral RNA synthesis with 5 nM and 10 nM (up to 50% for both doses) (**Fig. 2B**). We next evaluated whether VIP and PACAP could also be able to restrict virus production in pulmonary cells, one of the major targets of SARS-CoV-2. We found that VIP reduced viral replication, reaching up to 50% and 40% inhibition with 1 nM and 5 nM, respectively (**Fig. 2C and Supp Fig. 1A**). PACAP also diminished virus production up to 40% and 50% at concentrations equivalent to 10 nM and 50 nM (**Fig. 2D and Supp Fig. 1B**). In parallel, VIP and PACAP protected monocytes and Calu-3 cells from SARS-CoV-2-mediated cytopathic effect, as measured by LDH activity in supernatants (**Fig. 2E and 2F**). Overall, these results show that cells exposed to VIP or PACAP present decreased viral output and resistance to damages induced by the infection.

**Figure 2.**
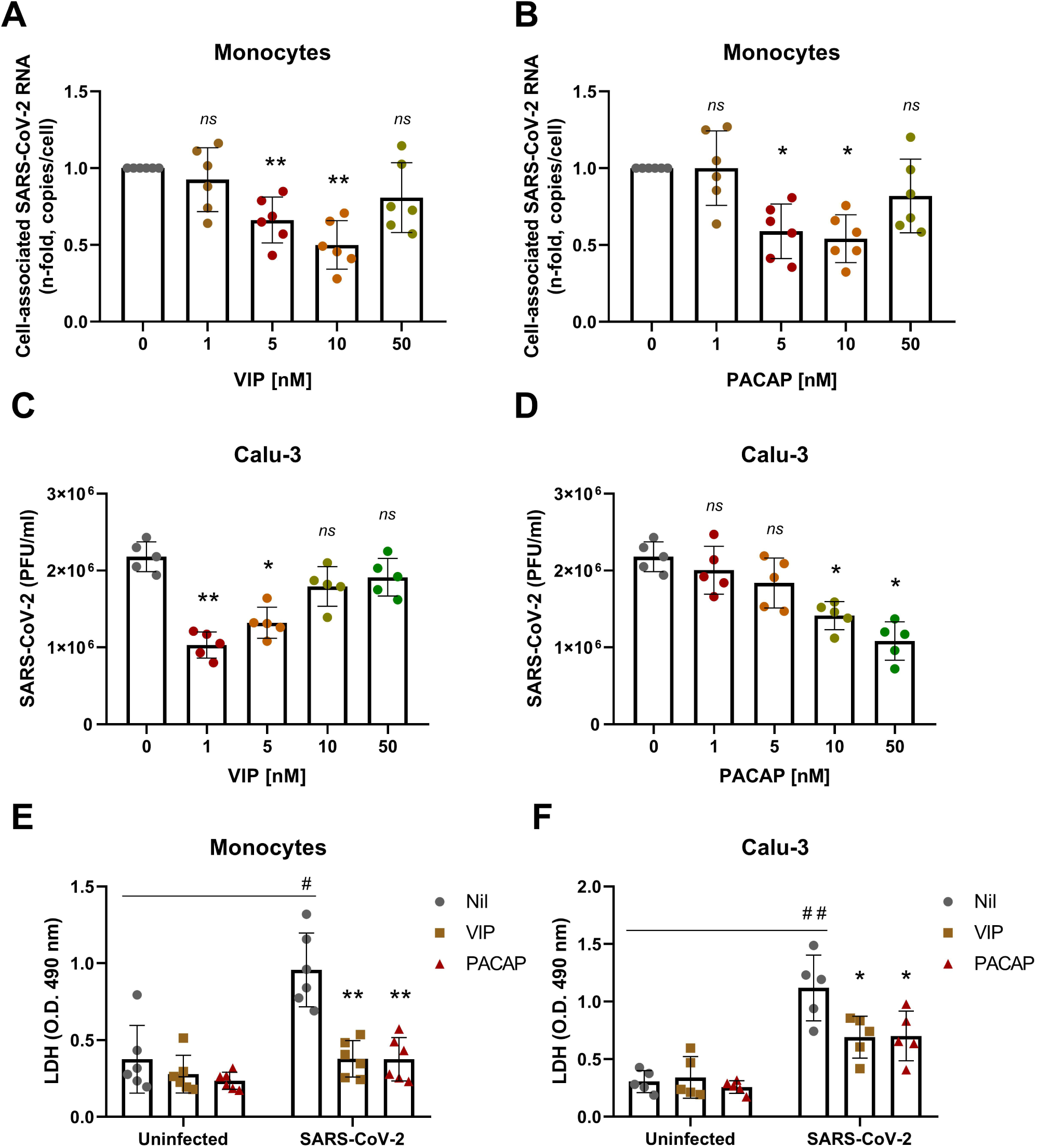
VIP and PACAP reduce SARS-CoV-2 RNA synthesis in human primary monocytes and viral replication in pulmonary cells, protecting them from virus-mediated cytopathic effects. Monocytes (**A, B**) and Calu-3 cells (**C, D**) were exposed (overnight) or not to the indicated concentrations of VIP (**A, C**) or PACAP (**B, D**). Culture medium was removed and then cells were infected with SARS-CoV-2. After infection, viral input was removed and cells were washed, then re-exposed to the neuropeptides. Viral RNA synthesis was evaluated by qPCR in monocytes 24 hours after infection. In Calu-3 cells, supernatants were collected at 48 hours after infection, and viral replication was evaluated by quantifying PFUs in Vero E6 plaque assays. Cellular viability was analyzed by measuring LDH release in the supernatants of uninfected or SARS-CoV-2-infected monocytes (**E**) treated or not with VIP or PACAP (10 nM), and Calu-3 cells (**F**) treated or not with VIP (1 nM) or PACAP (50 nM). Data in (**A, B**) are shown normalized to infected cells kept only with culture medium, and in (**C, D, E, F**) represent means ± SD of absolute values. \**p* ≤ .05; \*\**p* ≤ .01; (A, B, E) n=6; (C, D, F) n=5. Each dot represents an independent assay with three replicates.

### Receptor contribution for the VIP and PACAP mediated inhibition of SARS-CoV-2 replication

The different optimal concentrations of VIP and PACAP to reduce SARS-CoV-2 replication in Calu-3 cells might be explained by the relative abundance of the neuropeptide receptors, since it has been shown that these cells express only VPAC1 [43]. However, all three receptors are reported to be expressed in lungs, with some studies showing that VPAC1 levels are higher than VPAC2 or PAC1 (to which PACAP binds with higher affinity than to VPAC1 and VPAC2 [22,44–46]). With that in mind, we evaluated the role of the individual receptors in the neuropeptide-mediated inhibition of SARS-CoV-2 in both cells. To this end, monocytes were treated with specific agonists to VPAC1, VPAC2 and PAC1 (Ala-VIP, Bay 55-9837 and Maxadilan, respectively), and then infected with SARS-CoV-2. Activation of VPAC1 at 1 nM, 5 nM and 10 nM, and of VPAC2 at 1 nM, significantly reduced the SARS-CoV-2 genome replication (**Fig. 3A**). We also verified that VPAC1 is the main receptor involved the inhibition of SARS-CoV-2 in Calu-3 cells, resembling the level of inhibition achieved with VIP, while the exposure to a VPAC2 agonist resulted in a more modest inhibition (**Fig. 3B**). The stimulus with a PAC1 agonist had no effect on viral replication (**Fig. 3A and 3B**). As a whole, these findings suggest that VPAC1 receptor is the main contributor for the VIP- and PACAP-mediated SARS-CoV-2 inhibition in monocytes and Calu-3 cells, and that activation of this receptor can lead to a diminished viral replication similar to that induced by the own neuropeptides.

**Figure 3.**
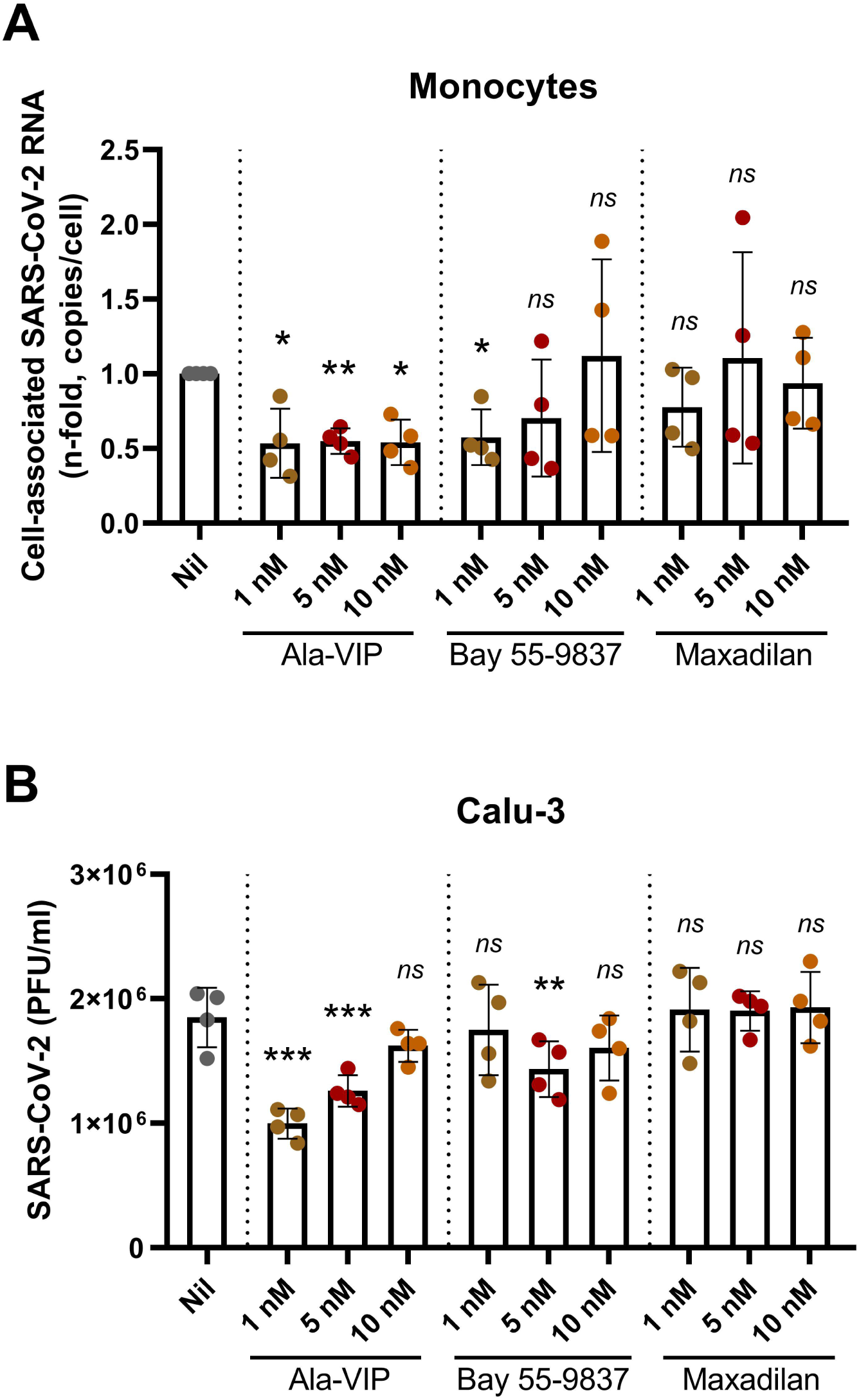
Receptor contribution for the VIP and PACAP mediated inhibition of SARS-CoV-2 replication. Monocytes (**A**) and Calu-3 cells (**B**) were treated (overnight) or not with agonists for VIP and PACAP receptors, as indicated, at different concentrations. Culture medium was removed and then cells were infected with SARS-CoV-2. After infection, viral input was removed, and cells were washed, and then re-exposed to the receptor agonists. Viral RNA synthesis was evaluated by qPCR in monocytes 24 hours after infection. In Calu-3 cells, supernatants were collected at 48 hours after infection, and viral replication was evaluated by quantifying PFUs in Vero E6 plaque assays. Data in (**A**) are shown normalized to infected cells kept only with culture medium, and in (**B**) represents means ± SD of absolute values. \**p* ≤ .05; \*\**p* ≤ .01; \*\*\**p* ≤ .001; (A, B) n=4. Each dot represents an independent assay with three replicates.

### VIP and PACAP reduce the production of proinflammatory cytokines by SARS-CoV-2-infected monocytes and Calu-3 cells

Controlling the production of proinflammatory cytokines may be critical for reducing SARS-CoV-2 replication and limiting tissue damages, and based on evidence that VIP and PACAP can regulate the inflammatory response [27,47], we next evaluated whether both neuropeptides could attenuate the production of proinflammatory mediators by SARS-CoV-2-infected monocytes or lung epithelial cells. As shown in **Fig. 4A**, SARS-CoV-2-infected monocytes produced large amounts of the proinflammatory mediators IL-6, IL-8, TNF and MIF relative to uninfected cells (15, 4, 12 and 18 times more, respectively). In contrast, the treatment of SARS-CoV-2-infected monocytes with either neuropeptide reduced to 66%, 50%, 66% and 50% the cellular production of IL-6, IL-8, TNF and MIF, respectively. Furthermore, VIP and PACAP reverted by approximately the same degree the release of IL-6 and IL-8 by Calu-3 cells (**Fig. 4B**), implying that VIP and PACAP may offer a critical protection to inflamed lungs affected by SARS-CoV-2 replication. Because proinflammatory cytokines may favor SARS-CoV-2 replication, which, in turn, can amplify the cellular synthesis of these mediators, these findings may support our assumption that VIP and PACAP offer tissue protection by inhibiting virus replication and regulating the boost of cytokine production.

**Figure 4.**
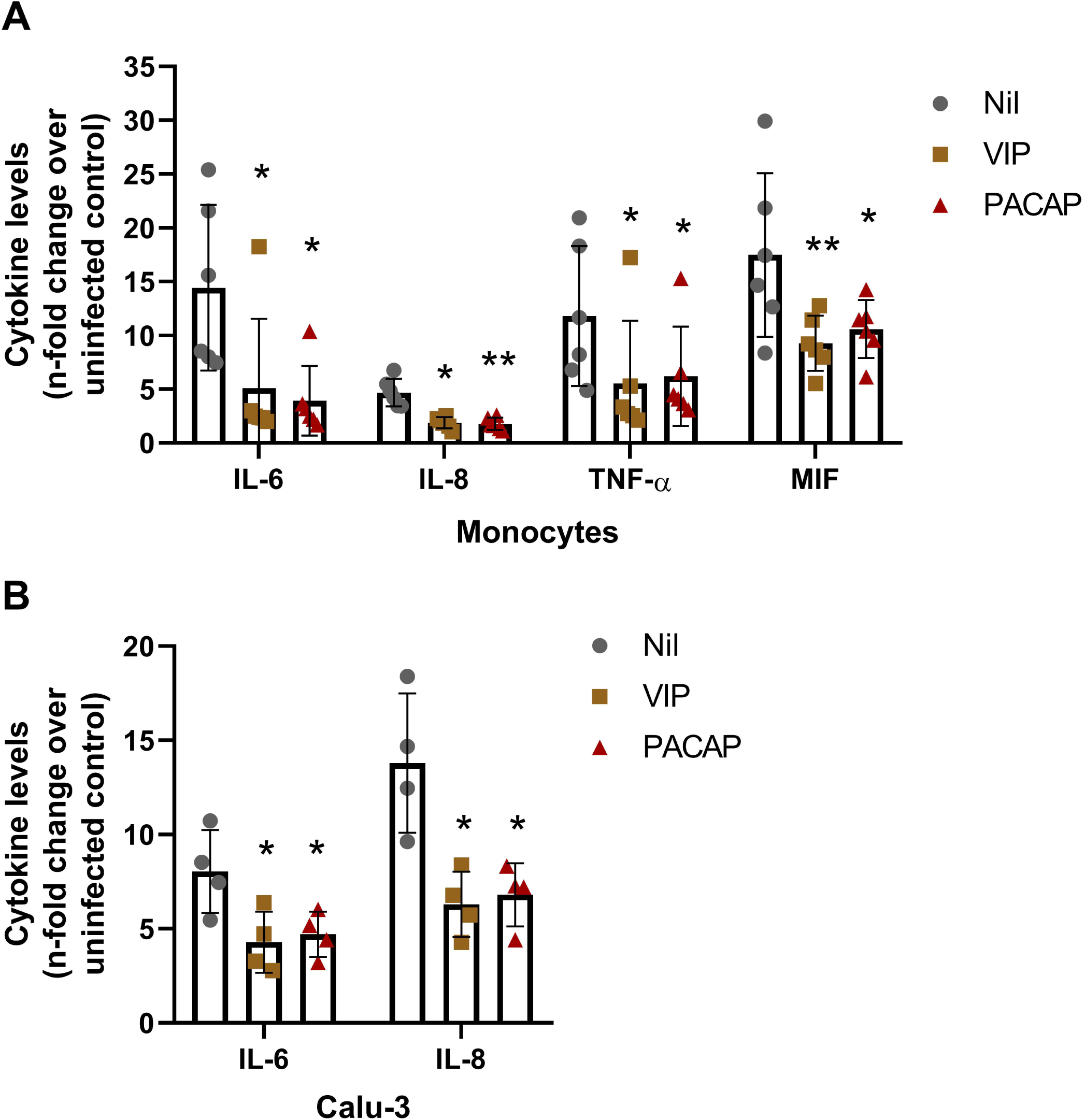
VIP and PACAP reduce the production of proinflammatory mediators by SARS-CoV-2-infected monocytes and Calu-3 cells. Monocytes (**A**) and Calu-3 cells (**B**) were treated (overnight) or not with VIP or PACAP (10 nM each for monocytes, 1 nM of VIP or 50 nM of PACAP for Calu-3 cells). Culture medium was removed and then cells were infected with SARS-CoV-2. After infection, viral input was removed and cells were washed, and then re-exposed to the neuropeptides. The levels of IL-6, IL-8, TNF-α and MIF were measured in culture supernatants of monocytes after 24 hours (**A**), and of IL-6 and IL-8 after 48 hours for Calu-3 cells (**B**), by ELISA. Data represent means ± SD. \**p* ≤ .05; \*\**p* ≤ .01; (A) n=6; (B) n=4. Each dot represents an independent assay with three replicates.

### VIP and PACAP regulate the activation of transcription factors in SARS-CoV-2-infected monocytes

Given that the transcription factor NF-kB is critically involved in the cellular production of inflammatory mediators [48], and our own findings showing that VIP and PACAP can inhibit its activation in HIV-1-infected macrophages [49], we investigated whether both neuropeptides would exert this same effect on SARS-CoV-2-infected monocytes. We found that activated NF-kB is up-modulated in infected cells (as measured by the increased amount of phosphorylated NF-kBp65 subunit), and that VIP and PACAP were able to reduce NF-KBp65 phosphorylation (**Fig. 5A**). Following, we analyzed the effects of both neuropeptides on the activation of CREB, a transcription factor induced by several GPCR ligands, including VIP and PACAP [50], and also involved in the induction of anti-inflammatory cytokines [51,52]. CREB and NF-kB share the CREB-binding protein/p300 (CBP/p300 protein) as a cofactor, and CREB activation results in the inhibition of NF-kB [53]. We found that activation of CREB was diminished in SARS-CoV-2-infected monocytes (**Fig. 5B**), a result coherent with NF-kB activation in the same cells. Consistent with this finding, VIP and PACAP promoted CREB activation (as measured by increase of CREB phosphorylation) in those infected monocytes, a result matching the inhibition of NF-kB and the reduction of cellular production of proinflammatory cytokines. We also evaluated in SARS-CoV-2-infected monocytes the expression of the active form of SREBP-1 and SREBP-2, transcription factors that also interact with CBP/p300 [54], and are crucial for the replication of several viruses, including coronaviruses [55–57]. In fact, we and other authors reported that SARS-CoV-2 infection promotes the activation of SREBP, and that this activation is associated with enhanced viral replication [58,59] and COVID-19 disease severity [60]. We detected that the levels of both isoforms of SREBP in active state are increased in SARS-CoV-2-infected monocytes and that VIP or PACAP treatment prevented this augmentation, lowering them to the same basal levels found in uninfected monocytes (**Fig. 5C and 5D, Supp. Fig. 2**).

**Figure 5.**
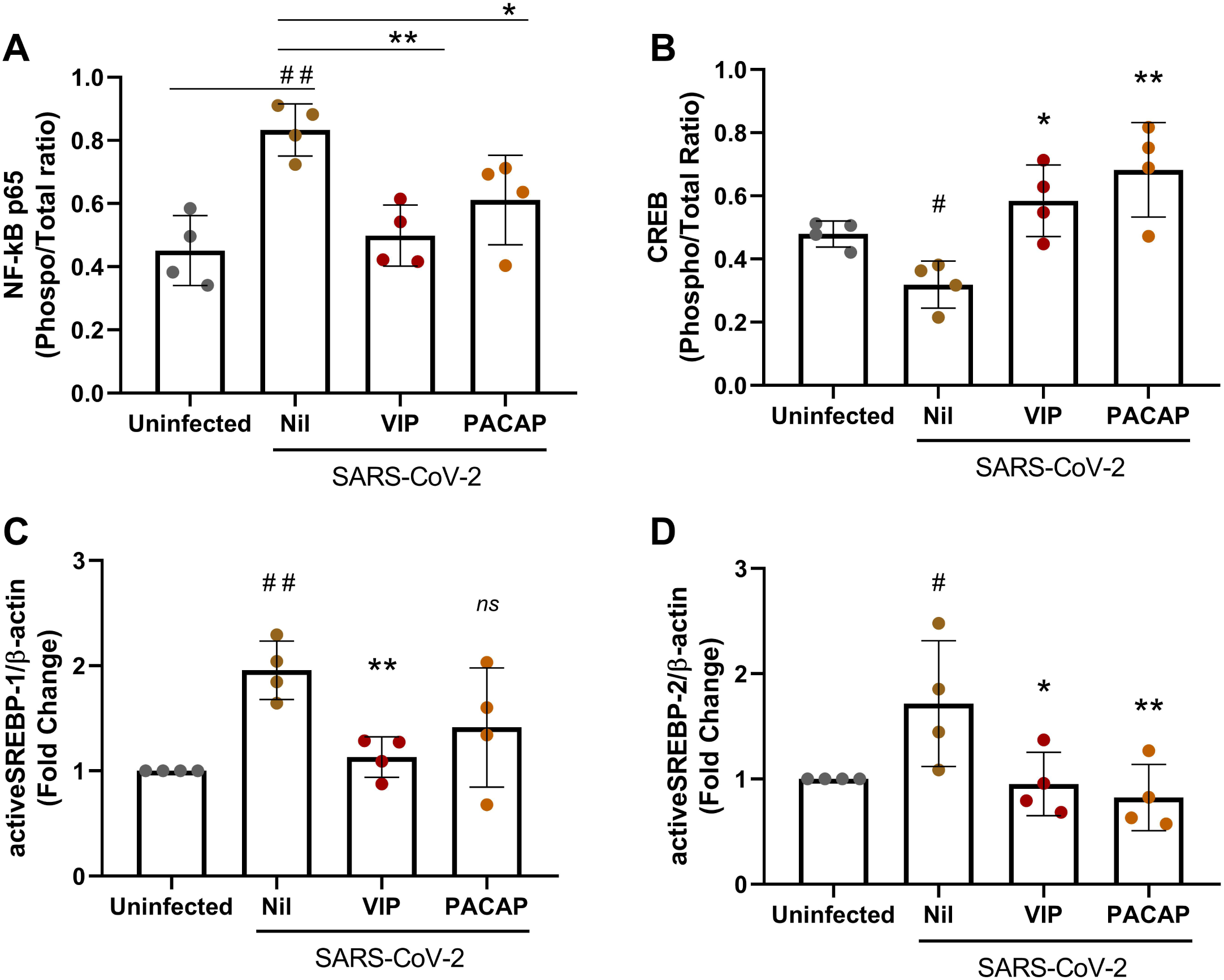
VIP and PACAP regulate the activation of transcription factors in SARS-CoV-2-infected monocytes. Monocytes were treated (overnight) or not with to VIP or PACAP (10 nM), culture medium was removed and then cells were infected with SARS-CoV-2. After infection, viral input was removed, and cells were washed and then re-exposed to the neuropeptides. After 24 hours, cells were lysed and the ratios between phosphoNF-kBp65 and total NF-kBp65 (**A**), phosphoCREB and total CREB (**B**), active SREBP-1 and β-actin (**C**), and active SREBP-2 and β-actin (**D**) were quantified by ELISA (**A, B**) or by western blot (**C, D**) in the cell lysates. Data represent means ± SD. \**p* ≤ .05; \*\**p* ≤ .01; \*\*\**p* ≤ .001; (A, B) n=3; (C, D) n=4. Each dot represents an independent assay with three replicates.

### Inhibition of NF-kB and SREBP in monocytes reduces SARS-CoV-2 RNA synthesis, production of proinflammatory mediators and protects the cells from virus-mediated cytopathic effects

To directly connect these latter findings with viral replication and production of pro-inflammatory mediators, we treated SARS-CoV-2-infected monocytes with pharmacological inhibitors of NF-KB (Bay 11-7082) or SREBP (AM580) [55,59], together or not with VIP and PACAP. We found that the sole inhibition of SREBP decreased viral RNA synthesis and production of TNF-α and IL-6, and reduced cell death, measurements that were all amplified when the inhibitors were associated with either neuropeptide (Fig. 6, A-D). Except for viral RNA synthesis, the sole inhibition of NF-KB, or in combination with VIP or PACAP, produced similar results (Fig. 6, A-D). Importantly, the protecting effects mediated by VIP or PACAP alone were identical to those seen when the signaling pathways triggered by NF-KB or SREBP activation were specifically inhibited.

**Figure 6.**
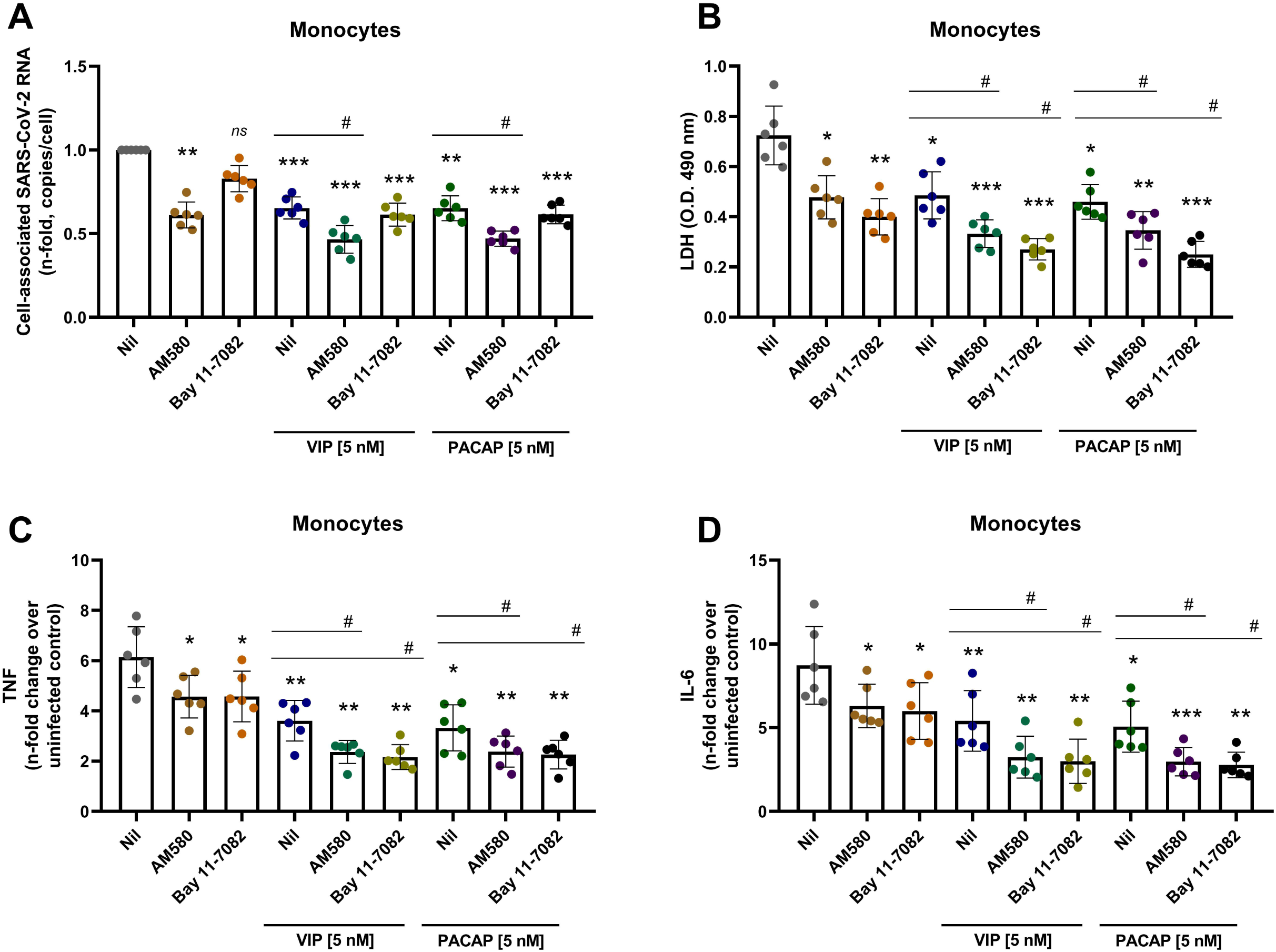
Inhibition of NF-kB and SREBP in monocytes reduces SARS-CoV-2 RNA synthesis, production of proinflammatory mediators and protects the cells from virus-mediated cytopathic effects. Monocytes were treated (overnight) or not with to VIP or PACAP (5 nM), culture medium was removed and then cells were infected with SARS-CoV-2. After infection, viral input was removed, and cells were washed and then re-exposed to the neuropeptides associated or not with inhibitors of SREBP (AM580, 10 uM) or NF-kB (Bay 11-7082, 10 uM). Viral RNA synthesis (A), cellular viability (B) and levels of TNF-α and IL-6 **(C, D)** were evaluated by qPCR, ELISA and LDH release, respectively, in the culture supernatants 24 hours after infection. Data in (**A**) are shown normalized to infected cells kept only with culture medium, and in (B, C, D) represent means ± SD of absolute values. \**p* ≤ .05; \*\**p* ≤ .01; \*\*\**p* ≤ .001; (A - D) n=6. Each dot represents an independent assay with three replicates.

Together, our data suggest that the restriction of SARS-CoV-2 replication in monocytes and in pulmonary cells by VIP and PACAP can be the outcome of the intrinsic modulation of inflammatory mediators and transcription factors that are involved directly and indirectly with the viral replication. Considering that VIP and PACAP regulate inflammatory reactions, it is possible that their increased circulating amounts reflect a counter-regulatory effect elicited by the dysregulated immune response typical of the more severe clinical status of COVID-19 patients. Since SARS-CoV-2-induced NF-kB and SREBP activation are key events involved in the elevated production of proinflammatory cytokines in COVID-19 [58,60,61], the inhibition of these transcription factors, associated with the reduction of proinflammatory cytokines and with the decrease of viral replication by VIP and PACAP, strengthen the potential of these neuropeptides as possible therapeutical candidates for COVID-19.

## Discussion

In this work, we identified that the plasma levels of the neuropeptide VIP are elevated in patients with severe forms of COVID-19, correlating with viral load, associated with reduced inflammation, and that the elevated VIP levels at ICU admission predicted patients’ favorable outcome, including association with patient survival. In in vitro SARS-CoV-2-infected monocytes and epithelial lung cells, the neuropeptides VIP and PACAP, endogenous molecules presenting anti-inflammatory properties, reduced the exacerbated synthesis of proinflammatory mediators, coupled with the inhibition of SARS-CoV-2 replication. Our findings support and encourage clinical trials with VIP in COVID-19 patients, which are in progress with intravenous [38] and inhaled [39,40] formulations and are expected to be full disclosed this year and next. An initial release of the data, as preprint, shows an increase in survival rates and reduction of IL-6 levels on those who received intravenous Aviptadil (VIP) [62]. Our present data may substantiate additional larger trials with VIP, an overlooked molecule associated with antiviral, anti-inflammatory and enhanced survival activities.

Both neuropeptides regulate the inflammatory response due to their ability to decrease the production of proinflammatory mediators, and to elicit the production of anti-inflammatory molecules. Given that VIP and PACAP and their receptors are systemically distributed, including lungs [22,63], brain, and gut, we believe that the anti-SARS-CoV-2 effects of both neuropeptides would not be restricted to the respiratory tract, as shown by many studies in other chronic inflammatory illnesses.

VIP and PACAP decreased SARS-CoV-2 genome replication in monocytes, while protecting them from virus-induced cytopathicity. By diminishing the intracellular levels of viral RNA and other viral molecules, VIP and PACAP could prevent the cell death by pyroptosis, which has been described as one of the main causes of cell damage during SARS-CoV-2 infection [13,14]. VIP and PACAP also diminished the production of the proinflammatory cytokines IL-6, IL-8, TNF-α and MIF by these cells, in agreement with the reported ability of these neuropeptides to regulate the inflammatory response [24–27,64]. We found similar results with lung epithelial cells, supporting that VIP and PACAP may offer a critical protection to inflamed lungs affected by SARS-CoV-2 replication. It is possible that the higher amounts of VIP in patients with severe forms of infection may reflect a counter-regulatory feedback elicited by the dysregulated immune response of these patients.

We detected that the transcription factor CREB, which can act as a negative regulator of NF-kB [65,66], is down-regulated in SARS-CoV-2-infected monocytes, in opposition to NF-kB activation in the same cells, and that VIP and PACAP reversed both phenomenon in infected monocytes. In some models [67–72], CREB activation is related to induction of anti-inflammatory cytokines concomitant with reduction of pro-inflammatory molecules and through competition with NF-kB by their shared co-activator protein CBP/p300 [51,65,66,72]. CREB activation is also involved with the anti-apoptotic response in monocytes and macrophages, during differentiation and inflammatory stimuli [73,74]. The imbalance between CREB and NF-kB, either as a direct effect of infection by SARS-CoV-2 or a consequence of exposure of bystander cells to viral products and inflammatory molecules, could be an important target for inhibition of SARS-CoV-2 deleterious effects, at least in monocytes and probably also in lung cells, as a similar imbalance between CREB and NF-kB was observed in an acute inflammatory pulmonary condition [53].

Induction of SREBP activity by SARS-CoV-2 was consistent with data showing its increase and association with COVID-19 severity in patients [60]. SREBP1 regulates the expression of genes of fatty acid biosynthesis, whilst SREBP2 regulates genes involved in cholesterol biosynthesis, intracellular lipid movement and lipoprotein import [75]. While crucial for metabolic homeostasis, both transcription factors are involved in pathologies when misbalanced or overactivated [75], and several viruses are reported to induce their activation, as the up-regulation of host lipid biosynthesis is a requirement for their optimal replication [55–57]. As reported by our group [58] and others authors [59], recently reported that SARS-CoV-2 activates SREBP-1 and other pathways of lipid metabolism in human cells, and that lipid droplets enhance viral replication and production of inflammatory mediators. Similar to NF-kB and CREB, the association of SREBPs with CBP/p300 [54] makes its function susceptible to the availability of this co-factor, which abundance can be low or high depending on the state of activation of NF-kB and CREB. Thus, the modulation of each one of these factors by VIP and PACAP can reflect a fine tuning of the transcriptional regulation of metabolic and inflammatory pathways, which in turn can affect the replication of SARS-CoV-2. Our results with inhibitors of SREBPs and NF-KB, used alone or in combination with either neuropeptide, provide further connection between the ability of VIP and PACAP to regulate the activity of these transcription factors and to control viral replication and production of pro-inflammatory mediators, as well as to reduce SARS-CoV-2-induced cell damages. The decline of viral genome replication and production of inflammatory cytokines secondary to SREBP blockage are in agreement with previous reports showing that this transcription factor is essential for replication of a broad range of viruses, including coronaviruses in Calu-3 cells [55–57,59] and contributes to cytokine storm in COVID-19 patients [60]. The diminished production of TNF-α and IL-6 in our assays due to NF-KB inhibition agrees with its well-known role to eliciting inflammatory responses. Overall, we believe that the protecting role of VIP and PACAP against SARS-CoV-2 infection in vitro can be explained, at least in part, by their ability to simultaneously regulate the signaling pathways elicited by these transcription factors. Our findings are summarized in the model presented in Fig. 7.

**Figure 7.**
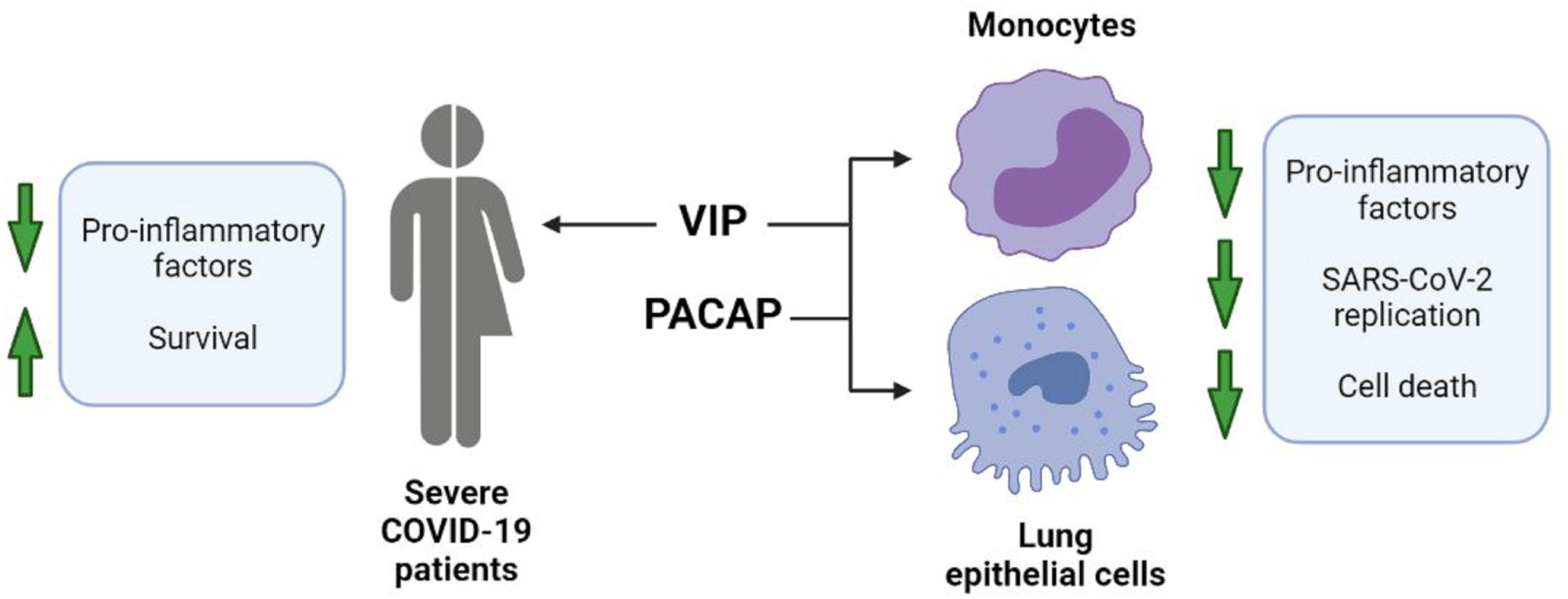
Graphical summary of study data. In severe COVID-19 patients, VIP plasma levels correlated with decreased inflammatory markers and survival. In in vitro assays with monocytes and lung epithelial cells, VIP and PACAP were found to decrease SARS-CoV-2 RNA synthesis (monocytes) and viral replication (lung epithelial cells). Both neuropeptides also reduced inflammatory factors and cell death of infected cells. Created with BioRender.com

Because VIP and PACAP signaling outcome is dependent of the combined action of the receptors activated by them (VIP and PACAP receptors can elicit cell signaling in homo and hetero dimers; 72), we evaluated whether they were involved in the final outcome analyzed. Our assays suggest that signaling through the receptors VPAC1 and VPAC2 contributed for VIP- and PACAP-mediated reduction of SARS-CoV-2 RNA synthesis in monocytes and viral production in Calu-3 cells, with VPAC1 activation alone being able to reproduce the SARS-CoV-2 inhibition promoted by the natural neuropeptides. The inhibition profile of SARS-CoV-2 by VIP and PACAP in Calu-3 cells may be biased regarding the expected action in the lungs, since Calu-3 cells appear to express only VPAC1 [43]. However, lung tissues, while reported to express high levels of VPAC1, also express VPAC2 and PAC1 [44,46], and, more specifically, VPAC2 mRNA was detected in airway epithelial, glandular, and immune cells of the lung [45]. Therefore, while the inhibition curve of SARS-CoV-2 by VIP and PACAP in Calu-3 cells points to different optimal doses than those obtained for monocytes, it is possible that in normal lung cells and tissue, VIP and PACAP could present a broader range of action in the inhibition of SARS-CoV-2. In fact, VIP and specific agonists for VPAC1 or VPAC2 have been proposed and tested for respiratory conditions, such as asthma [77–79], pulmonary arterial hypertension (PAH) [77,80,81] and chronic obstructive pulmonary disease (COPD) [77,78,82], demonstrating that the anti-inflammatory actions of VIP and PACAP can be achieved in lung tissues. Future studies should define which of these receptors would preferentially be activated by specific agonists to restrain SARS-CoV-2 replication in lungs or other sites. Also, as G-coupled receptors ligands, it is expected that VIP and PACAP curve profiles be subject to variation due receptor density in cell, receptor isoforms, and subtypes of associated G proteins. Those factors can influence the threshold and outcome of activation, and have been described for a variety of G-coupled receptors, including VIP/PACAP receptors [83– 85]. Together with the possible differences of receptor expression and self-regulatory characteristics of GPCRs, a third regulation level of VIP and PACAP action on pulmonary cells can be achieved by the activity of proteases and peptidases, as lungs are described to express high levels of several of them in both normal and pathological conditions [86–88]. Some of these peptidases could target VIP and PACAP, thus altering the ligand/receptor ratio and modulating the signaling pathways.

Since our results were obtained with the pandemic SARS-CoV-2 D614G, we cannot assure that the VIP and PACAP protective effects will also prevail against emerging variants. Nonetheless, given that both neuropeptides regulate cellular mechanisms, actions that might be independent of virus genotypes, we suppose that their protective effects will likely be replicated against SARS-CoV-2 variants.

As up to now the availability of antivirals specific to SARS-CoV-2 is limited, and that the hyper-inflammation may persist in COVID-19 patients even after the lowering of the viral load, the searching for compounds that target the aberrant production of proinflammatory cytokines and, simultaneously, the own viral replication, should be stimulated. Our present results showing that VIP and PACAP hold these two critical activities point these neuropeptides or their analogue molecules as potential therapeutic agents for COVID-19.

## Acknowledgments

We thank the Hemotherapy Service from Hospital Clementino Fraga Filho (Federal University of Rio de Janeiro, Brazil) for providing buffy-coats. Dr. Andre Sampaio from Farmanguinhos, platform RPT11M, and Dr. Lucio Mendes Cabral from Department of Drugs and Pharmaceutics, Faculty of Pharmacy, Federal University of Rio de Janeiro (UFRJ) are acknowledged for kindly donating the Calu-3 cell. The recombinant protein Maxadilan was kindly donated to us by Dr. Ethan A. Lerner (Department of Dermatology, Massachusetts General Hospital, MA, USA). The authors are thankful to Prof. Elvira M. Saraiva (Federal University of Rio de Janeiro, Brazil) for stimulating comments and invaluable suggestions. We pay tribute to Dr. Juliana de Meis, our cherished young colleague who died prematurely, leaving a significant legacy on knowledge of immunopathogenesis of parasitic diseases.

## Statement of Ethics

Experimental procedures involving human cells from healthy donors were performed with samples obtained after written informed consent and were approved by the Institutional Review Board (IRB) of the Oswaldo Cruz Institute/Fiocruz (Rio de Janeiro, RJ, Brazil) under the number 49971421.8.0000.5248. The National Review Board approved the study protocol (CONEP 30650420.4.1001.0008) for clinical samples, and informed consent was obtained from all participants or patients’ representatives.

## Author Contribution

Conceived the study: JRT, TMLS, DCBH; Designed the experiments: JRT, PTB, TMLS, DCBH; Performed the experiments: JRT, CQS, NFR, CRRP, CSF, SSGD, ACF, MM, VCS, LT, IGAQ, EDH, PK; Analyzed the data: JRT, PTB, IGAQ, EDH, PK, FAB, TMLS, DCBH; Wrote the paper: JRT, PTB, TMLS, DCBH. All authors reviewed and approved the manuscript.

## References

1 García LF. Immune Response, Inflammation, and the Clinical Spectrum of COVID-19. Front Immunol. 2020 Jun;11:1441.

2 Huang C, Wang Y, Li X, Ren L, Zhao J, Hu Y, et al. Clinical features of patients infected with 2019 novel coronavirus in Wuhan, China. Lancet. 2020 Feb;395(10223):497–506.

3 Chen G, Wu D, Guo W, Cao Y, Huang D, Wang H, et al. Clinical and immunological features of severe and moderate coronavirus disease 2019. J Clin Invest. 2020 Apr;130(5):2620–9.

4 Gupta A, Madhavan M V., Sehgal K, Nair N, Mahajan S, Sehrawat TS, et al. Extrapulmonary manifestations of COVID-19. Nat Med. 2020 Jul;26(7):1017–32.

5 Tang D, Comish P, Kang R. The hallmarks of COVID-19 disease. PLoS Pathog. 2020 May;16(5):e1008536.

6 Ragab D, Salah Eldin H, Taeimah M, Khattab R, Salem R. The COVID-19 Cytokine Storm; What We Know So Far. Front Immunol. 2020 Jun;11:1446.

7 Giamarellos-Bourboulis EJ, Netea MG, Rovina N, Akinosoglou K, Antoniadou A, Antonakos N, et al. Complex Immune Dysregulation in COVID-19 Patients with Severe Respiratory Failure. Cell Host Microbe. 2020 Jun;27(6):992-1000.e3.

8 Blanco-Melo D, Nilsson-Payant BE, Liu W-C, Uhl S, Hoagland D, Møller R, et al. Imbalanced Host Response to SARS-CoV-2 Drives Development of COVID-19. Cell. 2020 May;181(5):1036-1045.e9.

9 Veras FP, Pontelli MC, Silva CM, Toller-Kawahisa JE, de Lima M, Nascimento DC, et al. SARS-CoV-2-triggered neutrophil extracellular traps mediate COVID-19 pathology. J Exp Med. 2020 Dec;217(12):e20201129.

10 Radermecker C, Detrembleur N, Guiot J, Cavalier E, Henket M, D’Emal C, et al. Neutrophil extracellular traps infiltrate the lung airway, interstitial, and vascular compartments in severe COVID-19. J Exp Med. 2020 Dec;217(12):e20201012.

11 Skendros P, Mitsios A, Chrysanthopoulou A, Mastellos DC, Metallidis S, Rafailidis P, et al. Complement and tissue factor–enriched neutrophil extracellular traps are key drivers in COVID-19 immunothrombosis. J Clin Invest. 2020 Nov;130(11):6151–7.

12 Middleton EA, He XY, Denorme F, Campbell RA, Ng D, Salvatore SP, et al. Neutrophil extracellular traps contribute to immunothrombosis in COVID-19 acute respiratory distress syndrome. Blood. 2020 Sep;136(10):1169–79.

13 Rodrigues TS, de Sá KSG, Ishimoto AY, Becerra A, Oliveira S, Almeida L, et al. Inflammasomes are activated in response to SARS-cov-2 infection and are associated with COVID-19 severity in patients. J Exp Med. 2020 Mar;218(3):e20201707.

14 Ferreira AC, Soares VC, de Azevedo-Quintanilha IG, Dias S da SG, Fintelman-Rodrigues N, Sacramento CQ, et al. SARS-CoV-2 engages inflammasome and pyroptosis in human primary monocytes. Cell Death Discov. 2021 Jun;7(1):43.

15 Nicholls JM, Poon LLM, Lee KC, Ng WF, Lai ST, Leung CY, et al. Lung pathology of fatal severe acute respiratory syndrome. Lancet. 2003 May;361(9371):1773–8.

16 Gu J, Gong E, Zhang B, Zheng J, Gao Z, Zhong Y, et al. Multiple organ infection and the pathogenesis of SARS. J Exp Med. 2005 Aug;202(3):415–24.

17 Merad M, Martin JC. Pathological inflammation in patients with COVID-19: a key role for monocytes and macrophages. Nat Rev Immunol. 2020 Jun;20(6):355–62.

18 Schurink B, Roos E, Radonic T, Barbe E, Bouman CSC, de Boer HH, et al. Viral presence and immunopathology in patients with lethal COVID-19: a prospective autopsy cohort study. The Lancet Microbe. 2020 Nov;1(7):e290–9.

19 Chiang C-C, Korinek M, Cheng W-J, Hwang T-L. Targeting Neutrophils to Treat Acute Respiratory Distress Syndrome in Coronavirus Disease. Front Pharmacol. 2020 Oct;11:1576.

20 Zhou J, Chu H, Li C, Wong BHY, Cheng ZS, Poon VKM, et al. Active replication of middle east respiratory syndrome coronavirus and aberrant induction of inflammatory cytokines and chemokines in human macrophages: Implications for pathogenesis. J Infect Dis. 2014 Sep;209(9):1331–42.

21 Tynell J, Westenius V, Rönkkö E, Munster VJ, Melén K, Österlund P, et al. Middle east respiratory syndrome coronavirus shows poor replication but significant induction of antiviral responses in human monocyte-derived macrophages and dendritic cells. J Gen Virol. 2016 Feb;97(2):344–55.

22 Dickson L, Finlayson K. VPAC and PAC receptors: From ligands to function. Pharmacol Ther. 2009;121(3):294–316.

23 Ganea D, Delgado M. Vasoactive intestinal peptide (VIP) and pituitary adenylate cyclase-activating polypeptide (PACAP) as modulators of both innate and adaptive immunity. Crit Rev Oral Biol Med. 2002;13(3):229–37.

24 Kim WK, Kan Y, Ganea D, Hart RP, Gozes I, Jonakait GM. Vasoactive intestinal peptide and pituitary adenylyl cyclase-activating polypeptide inhibit tumor necrosis factor-α production in injured spinal cord and in activated microglia via a cAMP-dependent pathway. J Neurosci. 2000 May;20(10):3622–30.

25 Larocca L, Calafat M, Roca V, Franchi AM, Leiros CP. VIP limits LPS-induced nitric oxide production through IL-10 in NOD mice. Int Immunopharmacol. 2007;7(10):1343–9.

26 Gonzalez-Rey E, Delgado M. Vasoactive intestinal peptide inhibits cycloxygenase-2 expression in activated macrophages, microglia, and dendritic cells. Brain Behav Immun. 2008 Jan;22(1):35–41.

27 Delgado M, Munoz-Elias EJ, Gomariz RP, Ganea D. VIP and PACAP inhibit IL-12 production in LPS-stimulated macrophages. Subsequent effect on IFNγ synthesis by T cells. J Neuroimmunol. 1999 May;96(2):167–81.

28 Delgado M, Munoz-Elias EJ, Gomariz RP, Ganea D. Vasoactive intestinal peptide and pituitary adenylate cyclase-activating polypeptide prevent inducible nitric oxide synthase transcription in macrophages by inhibiting NF-kappa B and IFN regulatory factor 1 activation. J Immunol. 1999 Apr;162(8):4685–96.

29 Moody TW, Ito T, Osefo N, Jensen RT. VIP and PACAP: recent insights into their functions/roles in physiology and disease from molecular and genetic studies. Curr Opin Endocrinol Diabetes Obes. 2011;18(1):61–7.

30 Gonzalez-Rey E, Varela N, Chorny A, Delgado M. Therapeutical Approaches of Vasoactive Intestinal Peptide as a Pleiotropic Immunomodulator. Curr Pharm Des. 2007 Apr;13(11):1113–39.

31 Pozo D, Gonzalez-Rey E, Chorny A, Anderson P, Varela N, Delgado M. Tuning immune tolerance with vasoactive intestinal peptide: A new therapeutic approach for immune disorders. Peptides. 2007 Sep;28(9):1833–46.

32 Martínez C, Juarranz Y, Gutiérrez-Cañas I, Carrión M, Pérez-García S, Villanueva-Romero R, et al. A Clinical Approach for the Use of VIP Axis in Inflammatory and Autoimmune Diseases. Int J Mol Sci. 2019 Dec;21(1):65.

33 CDC. Real-time RT-PCR Primers and Probes for COVID-19 [Internet]. Centers Dis Control Prev. 2020 [cited 2020 Dec 4]. Available from: https://www.cdc.gov/coronavirus/2019-ncov/lab/rt-pcr-panel-primer-probes.html

34 Wu C, Chen X, Cai Y, Xia J, Zhou X, Xu S, et al. Risk Factors Associated With Acute Respiratory Distress Syndrome and Death in Patients With Coronavirus Disease 2019 Pneumonia in Wuhan, China. JAMA Intern Med. 2020 Jul;180(7):934.

35 Shi S, Qin M, Shen B, Cai Y, Liu T, Yang F, et al. Association of Cardiac Injury With Mortality in Hospitalized Patients With COVID-19 in Wuhan, China. JAMA Cardiol. 2020 Jul;5(7):802.

36 Guo T, Fan Y, Chen M, Wu X, Zhang L, He T, et al. Cardiovascular Implications of Fatal Outcomes of Patients With Coronavirus Disease 2019 (COVID-19). JAMA Cardiol. 2020 Jul;5(7):811.

37 Souza-Moreira L, Campos-Salinas J, Caro M, Gonzalez-Rey E. Neuropeptides as pleiotropic modulators of the immune response. Neuroendocrinology. 2011;94(2):89–100.

38 NCT0431169. Intravenous Aviptadil for Critical COVID-19 With Respiratory Failure [Internet]. [ClinicalTrials.gov] Bethesda Natl Libr Med (US). [cited 2020 Jul 21]. Available from: https://clinicaltrials.gov/ct2/show/NCT04311697

39 NCT04536350. Inhaled Aviptadil for the Treatment of Moderate and Severe COVID-19 [Internet]. [ClinicalTrials.gov] Bethesda Natl Libr Med (US). [cited 2020 Jul 25]. Available from: https://clinicaltrials.gov/ct2/show/NCT04360096

40 NCT04844580. A Clinical Study Evaluating Inhaled Aviptadil on COVID-19 [Internet]. [ClinicalTrials.gov] Bethesda Natl Libr Med (US). [cited 2021 Jul 1]. Available from: https://clinicaltrials.gov/ct2/show/NCT04844580

41 Boumaza A, Gay L, Mezouar S, Bestion E, Diallo AB, Michel M, et al. Monocytes and macrophages, targets of severe acute respiratory syndrome coronavirus 2: The clue for coronavirus disease 2019 immunoparalysis. J Infect Dis. 2021 Aug;224(3):395–406.

42 Zheng J, Wang Y, Li K, Meyerholz DK, Allamargot C, Perlman S. Severe Acute Respiratory Syndrome Coronavirus 2-Induced Immune Activation and Death of Monocyte-Derived Human Macrophages and Dendritic Cells. J Infect Dis. 2021 Mar;223(5):785–95.

43 Dérand R, Montoni A, Bulteau-Pignoux L, Janet T, Moreau B, Muller JM, et al. Activation of VPAC 1 receptors by VIP and PACAP-27 in human bronchial epithelial cells induces CFTR-dependent chloride secretion. Br J Pharmacol. 2004 Feb;141(4):698–708.

44 Busto R, Prieto JC, Bodega G, Zapatero J, Carrero I. Immunohistochemical localization and distribution of VIP/PACAP receptors in human lung. Peptides. 2000 Feb;21(2):265–9.

45 Groneberg DA, Hartmann P, Dinh QT, Fischer A. Expression and distribution of vasoactive intestinal polypeptide receptor VPAC2 mRNA in human airways. Lab Investig. 2001;81(5):749–55.

46 Fagerberg L, Hallstrom BM, Oksvold P, Kampf C, Djureinovic D, Odeberg J, et al. Analysis of the human tissue-specific expression by genome-wide integration of transcriptomics and antibody-based proteomics. Mol Cell Proteomics. 2014 Feb;13(2):397–406.

47 Delgado M, Garrido E, Martinez C, Leceta J, Gomariz RP. Vasoactive intestinal peptide and pituitary adenylate cyclase-activating polypeptides (PACAP27) and PACAP38) protect CD4+CD8+ thymocytes from glucocorticoid-induced apoptosis. Blood. 1996;87(12):5152–61.

48 Liu T, Zhang L, Joo D, Sun S-C. NF-κB signaling in inflammation. Signal Transduct Target Ther. 2017 Dec;2(1):17023.

49 Temerozo JR, de Azevedo SSD, Insuela DBR, Vieira RC, Ferreira PLC, Carvalho VF, et al. The Neuropeptides Vasoactive Intestinal Peptide and Pituitary Adenylate Cyclase-Activating Polypeptide Control HIV-1 Infection in Macrophages Through Activation of Protein Kinases A and C. Front Immunol. 2018 Jun;9(JUN):1336.

50 Schomerus C, Maronde E, Laedtke E, Korf HW. Vasoactive intestinal peptide (VIP) and pituitary adenylate cyclase-activating polypeptide (PACAP) induce phosphorylation of the transcription factor CREB in subpopulations of rat pinealocytes: immunocytochemical and immunochemical evidence. Cell Tissue Res. 1996;286(3):305–13.

51 Matt T. Transcriptional control of the inflammatory response: a role for the CREB-binding protein (CBP). Acta Med Austriaca. 2002;29(3):77–9.

52 Wen AY, Sakamoto KM, Miller LS. The Role of the Transcription Factor CREB in Immune Function. J Immunol. 2010 Dec;185(11):6413–9.

53 Shenkar R, Yum H-K, Arcaroli J, Kupfner J, Abraham E. Interactions between CBP, NF-κB, and CREB in the lungs after hemorrhage and endotoxemia. Am J Physiol Cell Mol Physiol. 2001 Aug;281(2):L418–26.

54 Toth JI, Datta S, Athanikar JN, Freedman LP, Osborne TF. Selective Coactivator Interactions in Gene Activation by SREBP-1a and −1c. Mol Cell Biol. 2004 Sep;24(18):8288–300.

55 Yuan S, Chu H, Chan JF-W, Ye Z-W, Wen L, Yan B, et al. SREBP-dependent lipidomic reprogramming as a broad-spectrum antiviral target. Nat Commun. 2019 Dec;10(1):120.

56 Cloherty APM, Olmstead AD, Ribeiro CMS, Jean F. Hijacking of Lipid Droplets by Hepatitis C, Dengue and Zika Viruses—From Viral Protein Moonlighting to Extracellular Release. Int J Mol Sci. 2020 Oct;21(21):7901.

57 Taylor HE, Linde ME, Khatua AK, Popik W, Hildreth JEK. Sterol Regulatory Element-Binding Protein 2 Couples HIV-1 Transcription to Cholesterol Homeostasis and T Cell Activation. J Virol. 2011 Aug;85(15):7699–709.

58 Dias S da SG, Soares VC, Ferreira AC, Sacramento CQ, Fintelman-Rodrigues N, Temerozo JR, et al. Lipid droplets fuel SARS-CoV-2 replication and production of inflammatory mediators. PLOS Pathog. 2020 Dec;16(12):e1009127.

59 Zhang S, Wang J, Cheng G. Protease cleavage of RNF20 facilitates coronavirus replication via stabilization of SREBP1. Proc Natl Acad Sci U S A. 2021 Sep;118(37). DOI: 10.1073/pnas.2107108118

60 Lee W, Ahn JH, Park HH, Kim HN, Kim H, Yoo Y, et al. COVID-19-activated SREBP2 disturbs cholesterol biosynthesis and leads to cytokine storm. Signal Transduct Target Ther. 2020 Dec;5(1):186.

61 Kircheis R, Haasbach E, Lueftenegger D, Heyken WT, Ocker M, Planz O. NF-κB Pathway as a Potential Target for Treatment of Critical Stage COVID-19 Patients. Front Immunol. 2020 Dec;11:3446.

62 Youssef JG, Lee R, Javitt J, Lavin P, Lenhardt R, Park DJ, et al. Increased Recovery and Survival in Patients With COVID-19 Respiratory Failure Following Treatment with Aviptadil: Report #1 of the ZYESAMI COVID-19 Research Group. SSRN Electron J. 2021 Aug DOI: 10.2139/ssrn.3830051

63 Said SI. The discovery of VIP: Initially looked for in the lung, isolated from intestine, and identified as a neuropeptide. Peptides. 2007 Sep;28(9):1620–1.

64 Delgado M, Munoz-Elias EJ, Martinez C, Gomariz RP, Ganea D. VIP and PACAP38 modulate cytokine and nitric oxide production in peritoneal. Ann N Y Acad Sci. 1999;897:401–14.

65 Parry GC, Mackman N. Role of cyclic AMP response element-binding protein in cyclic AMP inhibition of NF-kappaB-mediated transcription. J Immunol. 1997 Dec;159(11):5450–6.

66 Ollivier V, Parry GCN, Cobb RR, de Prost D, Mackman N. Elevated Cyclic AMP Inhibits NF-κB-mediated Transcription in Human Monocytic Cells and Endothelial Cells. J Biol Chem. 1996 Aug;271(34):20828–35.

67 Luan B, Yoon YS, Lay J Le, Kaestner KH, Hedrick S, Montminy M. CREB pathway links PGE2 signaling with macrophage polarization. Proc Natl Acad Sci U S A. 2015 Dec;112(51):15642–7.

68 Zhao L. Suppression of Proinflammatory Cytokines Interleukin-1 and Tumor Necrosis Factor-in Astrocytes by a V1 Vasopressin Receptor Agonist: A cAMP Response Element-Binding Protein-Dependent Mechanism. J Neurosci. 2004 Mar;24(9):2226–35.

69 Morris RHK, Tonks AJ, Jones KP, Ahluwalia MK, Thomas AW, Tonks A, et al. DPPC regulates COX-2 expression in monocytes via phosphorylation of CREB. Biochem Biophys Res Commun. 2008 May;370(1):174–8.

70 Ernst O, Glucksam-Galnoy Y, Bhatta B, Athamna M, Ben-Dror I, Glick Y, et al. Exclusive Temporal Stimulation of IL-10 Expression in LPS-Stimulated Mouse Macrophages by cAMP Inducers and Type I Interferons. Front Immunol. 2019 Aug;10:1788.

71 Barátki BL, Huber K, Sármay G, Matkó J, Kövesdi D. Inflammatory signal induced IL-10 production of marginal zone B-cells depends on CREB. Immunol Lett. 2019 Aug;212:14–21.

72 Avni D, Ernst O, Philosoph A, Zor T. Role of CREB in modulation of TNFα and IL-10 expression in LPS-stimulated RAW264.7 macrophages. Mol Immunol. 2010 Apr;47(7–8):1396–403.

73 Park JM, Greten FR, Wong A, Westrick RJ, Arthur JSC, Otsu K, et al. Signaling Pathways and Genes that Inhibit Pathogen-Induced Macrophage Apoptosis— CREB and NF-κB as Key Regulators. Immunity. 2005 Sep;23(3):319–29.

74 Cheng JC, Kinjo K, Judelson DR, Chang J, Wu WS, Schmid I, et al. CREB is a critical regulator of normal hematopoiesis and leukemogenesis. Blood. 2008 Feb;111(3):1182–92.

75 Shimano H, Sato R. SREBP-regulated lipid metabolism: convergent physiology — divergent pathophysiology. Nat Rev Endocrinol. 2017 Dec;13(12):710–30.

76 Harikumar KG, Morfis MM, Lisenbee CS, Sexton PM, Miller LJ. Constitutive formation of oligomeric complexes between family B G protein-coupled vasoactive intestinal polypeptide and secretin receptors. Mol Pharmacol. 2006;69(1):363–73.

77 Wu D, Lee D, Sung YK. Prospect of vasoactive intestinal peptide therapy for COPD/PAH and asthma: a review. Respir Res. 2011 Dec;12(1):45.

78 Onoue S, Yamada S, Yajima T. Bioactive analogues and drug delivery systems of vasoactive intestinal peptide (VIP) for the treatment of asthma/COPD. Peptides. 2007 Sep;28(9):1640–50.

79 Lindén A, Hansson L, Andersson A, Palmqvist M, Arvidsson P, Löfdahl CG, et al. Bronchodilation by an inhaled VPAC2 receptor agonist in patients with stable asthma. Thorax. 2003 Mar;58(3):217–21.

80 Hamidi SA, Lin RZ, Szema AM, Lyubsky S, Jiang YP, Said SI. VIP and endothelin receptor antagonist: An effective combination against experimental pulmonary arterial hypertension. Respir Res. 2011 Dec;12(1):141.

81 Hilaire RC, Murthya SN, Kadowitza PJ, Jeter JR. Role of VPAC1 and VPAC2 in VIP mediated inhibition of rat pulmonary artery and aortic smooth muscle cell proliferation. Peptides. 2010;31(8):1517–22.

82 Burian B, Angela S, Nadler B, Petkov V, Block LH. Inhaled Vasoactive Intestinal Peptide (VIP) improves the 6-minute walk test and quality of life in patients with COPD: The VIP/COPD-trial. Chest. 2006 Oct;130(4):121S.

83 Couvineau A, Laburthe M. VPAC receptors: structure, molecular pharmacology and interaction with accessory. Br J Pharmacol. 2012 May;166(1):42–50.

84 Gurevich V V., Gurevich E V. Biased GPCR signaling: Possible mechanisms and inherent limitations. Pharmacol Ther. 2020;211. DOI: 10.1016/j.pharmthera.2020.107540

85 Gether U. Uncovering molecular mechanisms involved in activation of G protein-coupled. Endocr Rev. 2000;21(1):90–113.

86 Van Der Velden VHJ, Wierenga-Wolf AF, Adriaansen-Soeting PWC, Overbeek SE, Möller GM, Hoogsteden HC, et al. Expression of aminopeptidase N and dipeptidyl peptidase IV in the healthy and asthmatic bronchus. Clin Exp Allergy. 1998;28(1):110–20.

87 Dreymueller D, Uhlig S, Ludwig A. ADAM-family metalloproteinases in lung inflammation: potential therapeutic targets. Am J Physiol Cell Mol Physiol. 2015 Feb;308(4):L325–43.

88 Bonda WLM, Iochmann S, Magnen M, Courty Y, Reverdiau P. Kallikrein-related peptidases in lung diseases. Biol Chem. 2018 Sep;399(9):959–71.

