## Supplemental Material for "VIP plasma levels associate with survival in severe COVID-19 patients, correlating with protective effects in SARS-CoV-2-infected cells"

[1]Laboratory on Thymus Research, Oswaldo Cruz Institute, Fiocruz, Rio de Janeiro, RJ, Brazil; [2]Laboratory of Immunopharmacology, Oswaldo Cruz Institute, Fiocruz, Rio de Janeiro, RJ, Brazil; [3]National Institute for Science and Technology on Innovation in Diseases of Neglected Populations (INCT/IDPN), Center for Technological Development in Health (CDTS), Fiocruz, Rio de Janeiro, RJ, Brazil; [4]Iguaçu University, Nova Iguaçu, RJ, Brazil; [5]Program of Immunology and Inflammation, Federal University of Rio de Janeiro, UFRJ, Rio de Janeiro, RJ, Brazil; [6]Laboratory of Immunothrombosis, Department of Biochemistry, Federal University of Juiz de Fora (UFJF), Juiz de Fora, Minas Gerais, Brazil; [7]Paulo Niemeyer State Brain Institute, Rio de Janeiro, RJ, Brazil; [8]D'Or Institute for Research and Education, Rio de Janeiro, RJ, Brazil; [9]Evandro Chagas National Institute of Infectious Diseases, Fiocruz, Rio de Janeiro, RJ, Brazil; [10]National Institute for Science and Technology on Neuroimmunomodulation, Oswaldo Cruz Institute, Fiocruz, Rio de Janeiro, RJ, Brazil.

**Short Title:** Protective effects of VIP and PACAP in SARS-CoV-2 infection

**Corresponding authors:** Jairo R. Temerozo and Dumith Chequer Bou-Habib
Laboratório de Pesquisas sobre o Timo, Instituto Oswaldo Cruz/Fiocruz
Av. Brasil, 4365 - Manguinhos - 21040-360 - Pav. Leônidas Deane – sala 510


**Key words:** SARS-CoV-2; COVID-19; VIP; PACAP; Neuropeptides

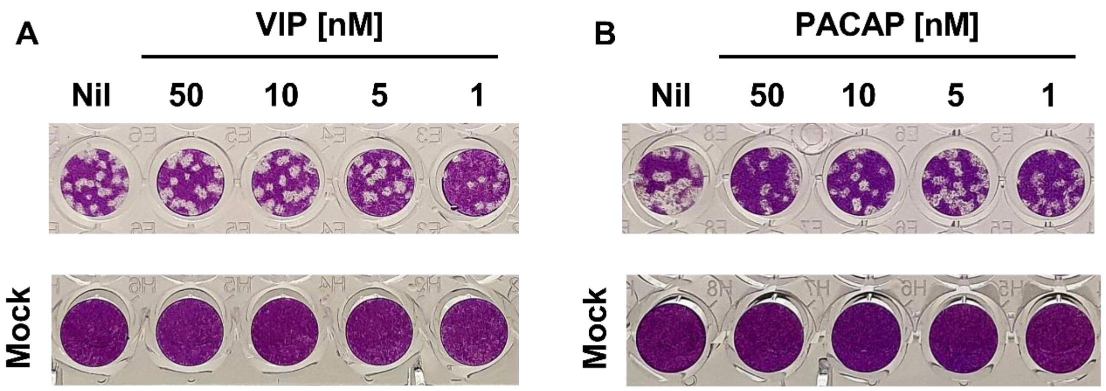


**Supplemental Figure 1**. **Representative images of PFU assays of SARS-CoV-2-infected Calu-3 cells exposed to VIP and PACAP.** Calu-3 cells were exposed (overnight) or not to the indicated concentrations of VIP (**A**) or PACAP (**B**). Culture medium was removed and then cells were infected with SARS-CoV-2. After infection, viral input was removed and cells were washed, then re-exposed to the neuropeptides. Supernatants were collected at 48 hours after infection, and viral replication was evaluated by quantifying PFUs in Vero E6 plaque assays.

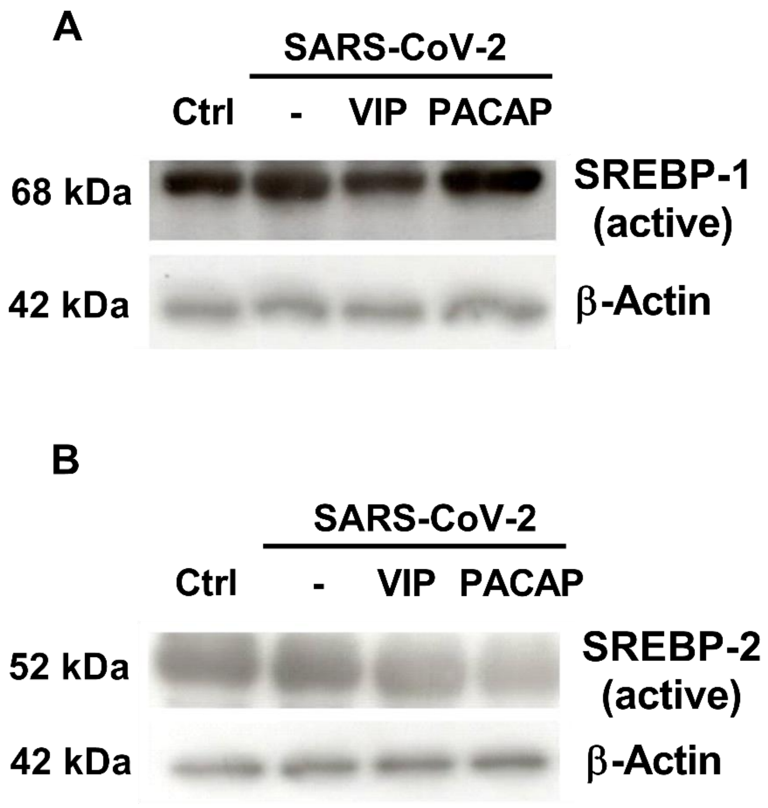


**Supplemental Figure 2**. **Representative blots of active-SREBP-1 and active-SREBP-2 in monocyte lysates.** Monocytes were treated (overnight) or not with to VIP or PACAP (10 nM) and then infected by SARS-CoV-2. After infection, cells were re-exposed to the neuropeptides. After 24 hours, cells were lysed and the ratios between active SREBP-1/β-actin (**A**), and active SREBP-2/β-actin (**B**) were quantified by western blot in the cell lysates. “Ctrl” indicates uninfected monocytes kept only with culture medium.

**Supplemental Table 1:** Characteristics of COVID-19 patients admitted to the ICU based on the requirement of noninvasive oxygen supplementation (moderate invasive mechanical) or ventilation (critical).

| Characteristics[1] | Moderate (N=5) | Critical (N=19) | p value[2] |
|---|---|---|---|
| Age, years | 38 (32 – 50) | 58 (48 – 67) | 0.0125 |
| Sex, male | 4 (80%) | 8 (42.1%) | 0.0611 |
| Respiratory support | | | |
| Oxygen supplementation | 5 (100%) | 19 (100%) | 1.0000 |
| Mechanical ventilation | 0 (0%) | 19 (100%) | <0.0001 |
| SAPS 3 | 31 (29 – 39) | 61 (58 – 75) | <0.0001 |
| $PaO_2/FiO_2$ ratio | 483 (372 – 519) | 139 (98 – 172) | 0.0021 |
| Vasopressor[3] | 0 (0%) | 10 (52.6%) | 0.0120 |
| Time from symptom onset to blood sample | 14 (7 – 15) | 13 (8 – 18) | 0.5767 |
| 28-day mortality | | | |
| **Comorbidities** | | | |
| Obesity | 2 (40%) | 3 (15.8%) | 0.3397 |
| Hypertension | 1 (20%) | 7 (36.8%) | 1.0000 |
| Diabetes | 2 (40%) | 7 (36.8%) | 1.0000 |
| Cancer | 0 (0%) | 3 (15.8%) | 0.5531 |
| Heart disease[3] | 0 (0%) | 2 (10.5%) | 1.0000 |
| **Presenting symptoms** | | | |
| Cough | 4 (80%) | 13 (68.4%) | 1.0000 |
| Fever | 4 (80%) | 15 (79%) | 1.0000 |
| Dyspnea | 5 (100%) | 15 (79%) | 0.6352 |
| Headache | 0 (0%) | 3 (15.8%) | 0.2977 |
| Anosmia | 3 (60%) | 5 (26.3%) | 0.4159 |
| **Laboratory findings on admission** | | | |
| Leukocytes, x 1000/μL | 98 (35 – 110) | 143 (122 – 192) | 0.0036 |
| Lymphocyte, cells/μL | 1269 (883 – 2343) | 1167 (518 – 1590) | 0.3412 |
| Monocytes, cells/μL | 495 (214 – 714) | 712 (563 – 908) | 0.1009 |
| Platelet count, x 1000/μL | 223 (157 – 320) | 205 (148 – 268) | 0.8375 |
| C Reactive Protein, mg/L[4] | 51 (17 – 204) | 208 (137 – 368) | 0.0518 |
| Fibrinogen, mg/dL[4] | 425 (368.4 – 572.6) | 545 (366 – 621) | 0.5392 |
| D-dimer, IU/mL[4] | 723 (285 – 4346) | 6235 (3337 – 14179) | 0.0089 |

[1]Numerical variables are represented as the median and the interquartile range, and qualitative variables are represented as the number and the percentage.
[2]Qualitative variables were compared using the two tailed Fisher exact test, and numerical variables using t test for parametric and Mann Whitney test for nonparametric distributions.
[3]Coronary artery disease or congestive heart failure.
[4]Reference values of C reactive Protein (0.00 – 1.00), Fibrinogen (238 – 498 mg/dL) and D-dimer (0 – 500 ng/mL).

**Supplemental Table 2:** Characteristics of COVID-19 patients admitted to ICU based on evolution to hospital discharge (survivors) or death (nonsurvivors).

| **Characteristics**[1] | **Survivors (N=11)** | **Nonsurvivors (N=13)** | **p value**[2] |
|---|---|---|---|
| Age, years | 47 (38 – 62) | 56 (50 – 73) | 0.0828 |
| Sex, male | 5 (36.7%) | 8 (61.5%) | 0.0529 |
| Respiratory support | | | |
| Oxygen supplementation | 5 (45.4%) | 13 (100%) | 0.0010 |
| Mechanical ventilation | 6 (54.5%) | 13 (100%) | 0.0046 |
| SAPS 3 | 55 (31 – 60) | 68 (59 – 78) | 0.0344 |
| $PaO_2/FiO_2$ ratio | 206 (132.5 – 427) | 123.5 (76.5 – 145) | 0.0558 |
| Vasopressor | 3 (27.3%) | 7 (53.8%) | 0.1519 |
| Time from symptom onset to blood sample | 13 (8 – 16) | 14 (7 – 18) | 0.2500 |
| **Comorbidities** | | | |
| Obesity | 3 (27.3%) | 2 (15.4%) | 0.3845 |
| Hypertension | 8 (72.7%) | 8 (61.5%) | 1.0000 |
| Diabetes | 3 (27.3%) | 5 (38.5%) | 1.0000 |
| Cancer | 2 (18.2%) | 1 (7.7%) | 1.0000 |
| Heart disease[3] | 1 (9.1%) | 1 (7.7%) | 0.5956 |
| **Presenting symptoms** | | | |
| Cough | 8 (72.7%) | 9 (69.2%) | 1.0000 |
| Fever | 9 (81.8%) | 10 (76.9%) | 0.6956 |
| Dyspnea | 9 (81.8%) | 11 (84.6%) | 0.9483 |
| Headache | 1 (9.1%) | 2 (15.4%) | 0.5815 |
| Anosmia | 5 (36.7%) | 3 (23.1%) | 0.6197 |
| **Laboratory findings on admission** | | | |
| Leukocytes, x 1000/µL | 106 (76 – 149) | 143 (122 – 251) | 0.0191 |
| Lymphocyte, cells/µL | 1470 (961 – 1596) | 934 (328 – 1647) | 0.2290 |
| Monocytes, cells/µL | 495 (448 – 742) | 738 (599 – 1005) | 0.0229 |
| Platelet count, x 1000/µL | 277 (161 – 347) | 194 (142 - 251) | 0.1641 |
| C Reactive Protein, mg/L[4] | 178 (51 – 336) | 199 (90 – 311) | 0.5538 |
| Fibrinogen, mg/dL[4] | 489 (366 – 581) | 545 (370 – 621) | 0.4072 |
| D-dimer, IU/mL[4] | 2499 (723 – 6235) | 7401 (3420 – 21706) | 0.0071 |

[1]Numerical variables are represented as the median and the interquartile range, and qualitative variables are represented as the number and the percentage.
[2]Qualitative variables were compared using the two tailed Fisher exact test, and numerical variables using t test for parametric and Mann Whitney test for nonparametric distributions.
[3]Coronary artery disease or congestive heart failure.
[4]Reference values of C reactive Protein (0.00 – 1.00), Fibrinogen (238 – 498 mg/dL) and D-dimer (0 – 500 ng/mL).
